# Stacked regressions and structured variance partitioning for interpretable brain maps

**DOI:** 10.1101/2023.04.23.537988

**Authors:** Ruogu Lin, Thomas Naselaris, Kendrick Kay, Leila Wehbe

## Abstract

Relating brain activity associated with a complex stimulus to different properties of that stimulus is a powerful approach for constructing functional brain maps. However, when stimuli are naturalistic, their properties are often correlated (e.g., visual and semantic features of natural images, or different layers of a convolutional neural network that are used as features of images). Correlated properties can act as confounders for each other and complicate the interpretability of brain maps, and can impact the robustness of statistical estimators. Here, we present an approach for brain mapping based on two proposed methods: *stacking* different encoding models and *structured variance partitioning*. Our stacking algorithm combines encoding models that each use as input a feature space that describes a different stimulus attribute. The algorithm learns to predict the activity of a voxel as a linear combination of the outputs of different encoding models. We show that the resulting combined model can predict held-out brain activity better or at least as well as the individual encoding models. Further, the weights of the linear combination are readily interpretable; they show the importance of each feature space for predicting a voxel. We then build on our stacking models to introduce structured variance partitioning, a new type of variance partitioning that takes into account the known relationships between features. Our approach constrains the size of the hypothesis space and allows us to ask targeted questions about the similarity between feature spaces and brain regions even in the presence of correlations between the feature spaces. We validate our approach in simulation, showcase its brain mapping potential on fMRI data, and release a Python package. Our methods can be useful for researchers interested in aligning brain activity with different layers of a neural network, or with other types of correlated feature spaces.

## Introduction

Over the past few years, the shift to a more complex, naturalistic experimental design has provided large amounts of experimental data [1, 2, 3, 4]. These data enable us to explore cognitive processes and map them on the brain computationally. Encoding models, a core class of brain mapping methods, focus on identifying the transformation from stimulus to brain activity [5]. They have been used to study multiple cognitive domains, such as visual, language, and auditory processing [5, 6, 7, 8, 9, 10, 11, 12, 13, 14, 15]. Linear encoding models are the most common, as they provide computational efficiency and interpretability. Recently, due to fast progress in artificial intelligence, encoding models have frequently been used in combination with feature spaces extracted from deep neural networks. Typically, these deep networks have many different components (for example, different layers in convolutional neural networks or transformer networks) [16, 17, 18, 13]. Researchers have also compared different networks [18, 19], or even the same layer of the same network architecture trained for different tasks [20]. This approach leads to important questions at the boundary of machine learning and neuroscience: how different are the representations at those layers and the functions they perform, and what does the mapping of the layers onto different “units” (voxels, electrophysiological sensors, etc.) of brain activity tell us about those units? Due to the number of layers and the correlation between their representations, it is hard to pick out which layer is most important for predicting brain activity, even when using linear models. Although encoding methods based on linear regression (typically ridge regression) have been shown to be useful for modeling functional magnetic resonance imaging (fMRI) data, they suffer from several complications. First, state-of-the-art encoding methods often use a single input feature space to describe the stimulus. This results in non-optimal encoding performance, since considering only one feature space might not capture all of the predictable signals. Second, some other approaches rely on combining multiple feature spaces by simply concatenating them. This is usually done in a setting in which a different penalty parameter is estimated at each voxel (volumetric pixel). As different feature spaces might predict different fractions of the variance of the activity in a voxel, choosing the same penalty parameter for all of them (1) does not give enough information to the model to treat features in a feature space as a group and (2) does not incorporate enough information about groups of features to estimate a different magnitude of weights for each group. To counteract this problem, Nunez-Elizalde et al. [21] proposed a banded-ridge setting in which each feature space can be given a different penalty parameter in a Tikhonov regression framework. This approach allows for a more informed and flexible model learning that can estimate the importance of feature spaces in the same voxel. The different penalties are estimated through a hyperparameter optimization setup that is quite computationally intensive, but has since been made more efficient through a new GPU implementation [22]. Another approach that combines information across feature spaces is the feature-weighted receptive field (fwRF) model [23]. fwRF encodes brain activity as a combination of multiple feature maps, constrained by a common spatial receptive field. Optimized by minimizing the least-squares cost for each voxel, it manages to recover voxel receptive field-like properties and tuning functions.

In this paper, we propose two methods that can be combined together for building more robust and easier to interpret brain maps. The first method uses an alternative algorithm (*stacked regression*) to combine individual feature spaces. The second method (*structured variance partitioning*) assesses the contribution of each feature space through a hypothesis testing scheme that incorporates knowledge about the relationships between feature spaces. To elaborate on the first method, we propose that a good way to combine the predictions from the individual feature spaces is to aim at optimizing their combined prediction performance, and a natural way to achieve this is via stacking. Wolpert [24] introduced stacking, an ensemble method that combines the output of multiple models to generate a new prediction. Breiman [25] proposed stacked regressions, a method for forming linear combinations of different regressors to improve prediction accuracy and determine the weights in the combination. Since its introduction in the 1990s, stacking has become a commonly used tool for combining classifiers. It is used in multiple scenarios [26, 27, 28, 29] and is successful in online competitions like Kaggle [30, 31].

Briefly, the stacked regressions approach we implement from Breiman [25] has two levels. The first level consists of the predictions made by different models. At the second level, the predictions are combined using a convex combination (a convex combination of *k* items *y*_1_, …, *y*_*k*_ is a linear combination 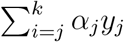 with weights *α*_*j*_ such that 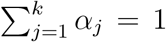 and 0 ≤ *α*_*j*_ ≤ 1 for *j* ∈ 1, …, *k*). The parameters of the combination *α*_*j*_ are learned through a quadratic optimization that minimizes the product of the residuals from different feature spaces (see methods). The effect of this procedure is that larger weights tend to be put on accurate predictors, while also balancing between predictors to select ones with less correlated errors over samples. If two good predictors have uncorrelated errors, they will make errors on different samples, and more samples will be well predicted by at least one of the predictors. Predictors with uncorrelated errors can thus complement each other to achieve better predictions.

We adapt the stacked regressions approach to the problem of building encoding models for the brain (see Figure 1). Crucially, we define the first level of our stacking procedure as consisting of linear regressors, each having a different stimulus feature space as input. At the second level, we learn the parameters *α*_*j*_ of a convex combination of first level predictors. The entire stacked model is estimated separately at each voxel. We show here using simulated data and real fMRI data from the Natural Scenes Dataset (NSD) [1] that stacking *k* predictors reliably leads to a model that is better or at least as good as the best of those *k* predictors. Further, we show that our stacking method provides robust and interpretable brain maps. After our model is learned, we can readily interpret the parameters of the convex combination: the 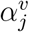 associated with feature space **x**_*j*_ at voxel *v* corresponds to the estimated importance of **x**_*j*_ for predicting voxel *v*.

**Figure 1:**
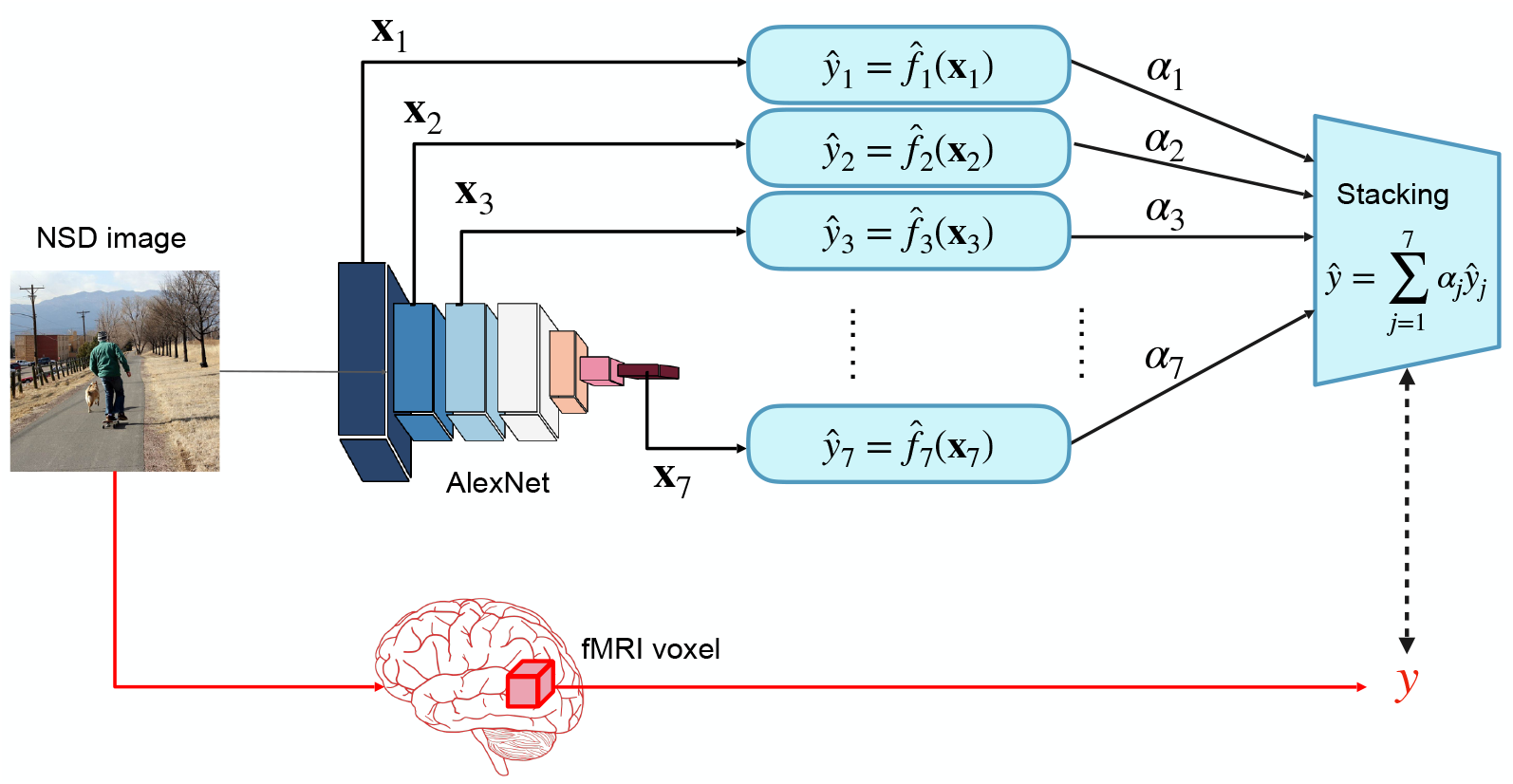
Schematic of our stacking approach. To build a stacked encoding model for a voxel, we start with multiple feature spaces **x**_1_, **x**_2_, …, **x**_*n*_ that describe stimulus properties. In this example, the fMRI recordings are from the Natural Scenes Dataset (NSD) [1], the stimulus is a set of natural scenes from the large COCO image database [32], and the features are the activations at different layers of AlexNet [33] corresponding to each image. We consider here only one voxel. We independently train regressors *f*_*j*_ for that voxel that each take as input only one feature space **x**_*j*_. Each 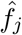 produces a prediction *ŷ*_*j*_ of the voxel’s activity. The parameters *α*_*j*_ of a convex combination (i.e., *α*_*j*_ ≥ 0 for all *j* and *α*_*j*_ = 1) are learned to produce an optimal combination of the predictions: *ŷ* = ∑_*j*_ *α*_*j*_*ŷ*_*j*_. We evaluate the predictions on held-out data.

The second method we propose, *structured variance partitioning*, uses knowledge about the interdependence of feature spaces to relate these feature spaces to brain regions. This approach builds on variance partitioning [34, 35, 36], a method used to determine the relative importance of sets of variables in regression models. Because of the correlation between feature spaces, variance partitioning is often used to capture the amount of variance that is predicted by information unique to one feature space or a group of feature spaces. This is often done by contrasting the variance predicted using all feature spaces, and the one predicted while excluding the (group of) feature space(s) of interest [35, 36]. Inherent to the assumptions of variance partitioning is that combining additional feature spaces should not decrease the prediction performance. However, we show in this paper that the typical way to combine feature spaces (concatenation) can often result in a decrease in performance due to statistical limitations (i.e., additional feature spaces places greater demands on data for accurate parameter estimation and may therefore lead to decreased performance on out-of-sample data), and that stacking is an efficient and useful way to perform variance partitioning in a way that preserves the required assumptions.

Crucially, we extend variance partitioning to make use of the experimenter’s knowledge of the relationship between feature spaces (such as the consecutive layers of a network, or other feature spaces that are assumed to relate to each other [37]) to perform more targeted hypothesis tests. In classical variance partitioning, if the modeling problem includes *M* feature spaces, fully investigating the problem might require performing variance partitioning an exponential number of times, which is computationally expensive and, more importantly, hard to interpret. For example, one might look at the unique variance explained by each of the *k* feature spaces separately (*k* variance partitioning calculations), or at the unique variance explained by all pairs of feature spaces (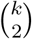 variance partitioning calculations), and so on. It should be evident that such a large number of computations, even if feasible, would be quite hard to make sense of. In many cases, we know exactly how feature spaces relate to one another (e.g., the successive layers of a convolutional neural network are functions of each other), and thus we can use these relationships to vastly reduce the number of variance partitioning analyses by incorporating this structure.

In the following sections, we explain the technical details of our proposed methods and show their behavior on simulated data and on real fMRI data. This paper is accompanied by a Python package that can be found at https://github.com/brainML/Stacking_Basics.

## Methods

In this section, we describe the setup of the stacking approach, the simulation experiments, and the real data used.

### Voxelwise encoding models

We denote the data we collected in fMRI experiments in the following way: Brain activity **y**^(*i*)^ of a subject are recorded at time points or trials *i* = 1, …, *n* while they are exposed to stimuli *s*^(*i*)^. Each brain measurement consists of many voxels, e.g. **y**^(*i*)^ = [*y*^1(*i*)^, *y*^2(*i*)^, …, *y*^*m*(*i*)^]. The goal of an encoding model is to find a function that relates each voxel to the stimulus: *y*^*v*(*i*)^ = *h*_*v*_ *s*^(*i*)^. To capture the relationship between *y*^*v*^ and *s*, it is common to use a feature space **x** that represents the properties of the stimulus [5]. Brain activity is then estimated as a function of the feature space: 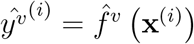. It is common for the function *f* to be linear and fit using a ridge penalty [5, 10, 38, 12, 11, 39].

### First level regressors

We are given tuples of brain activity and *k* corresponding stimulus feature spaces 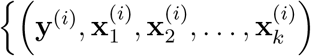 for *i* ∈ 1, …, *n*}, with the feature spaces describing different stimulus properties. We use ridge regression as our first-level regressor due to its stability, ease of computation and the fact that it is a common choice for encoding models. For each feature space **x**_*j*_, we train a separate encoding model 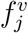 to predict the activity *y*^*v*^ parameter 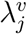, i.e. 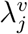 is *chosen independently for each feature space j and each voxel v* in a in voxel *v* as 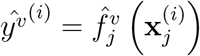. We train our first level ridge regressors with a voxelwise ridge nested cross-validation procedure (possible *λ*_*j,v*_ values range from 10^−6^ to 10^10^).

### Stacked Regression

We adapt the stacked regression method by Breiman [25] to build encoding models that combine multiple feature spaces. We have a set of encoding models *f*_1_, …, *f*_*k*_. The key change from the classical stacking approach is that we use linear regression models with different feature spaces as input, where usually in stacking, different types of predictors (e.g. linear and non-linear) are used with the same input features. For simplicity of notation, we consider here only one voxel, but the procedure below is applied at each voxel independently. Instead of selecting an individual encoding model from *f*_1_, …, *f*_*k*_, a more accurate predictor can be obtained by combining the encoding models such that:

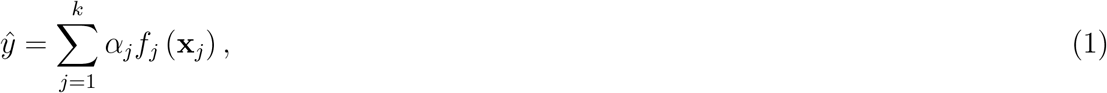

where the *α*_*j*_s are chosen according to a specific optimization problem. Given paired

*y*, (**x**_1_, **x**_2_, …, **x**_*k*_) and encoders *f*_*j*_ (**x**_*j*_) trained to predict *y* from **x**_*j*_, we want to solve:

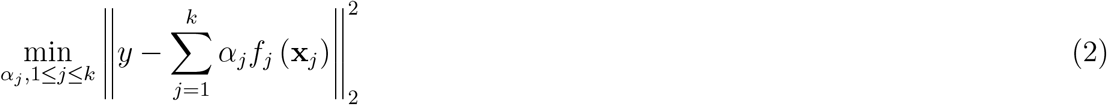

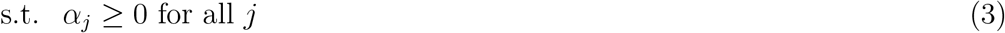

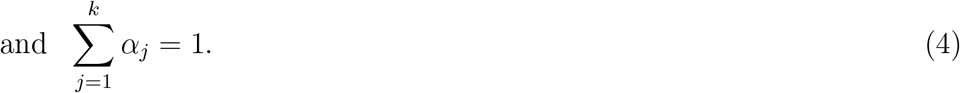

These constraints guarantee that the stacked prediction is a convex combination of the individual predictions. Note that

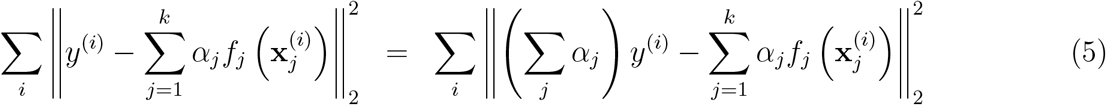

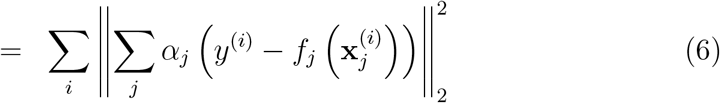

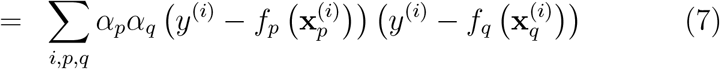

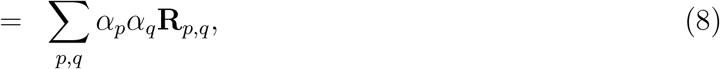

where *i* is the sample index and **R** is a matrix containing the residual products:

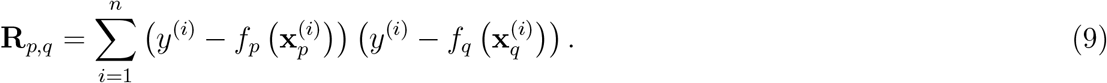

In keeping with Breiman [25], we select the parameters of the convex combination using the residuals of the first-level models estimated on held-out data. To make use of the entire training dataset, we use a nested cross-validation setting in which we hold out 80% of the training data, estimate all *f*_*j*_s on these held out data (via nested cross-validation), then produce predictions for the 20% held out data from all 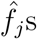 (see Figure 2 and Algorithm 1). We repeat this over all folds and combine all the predictions so that they have the original size as the training data. We compute the matrix **R** using these predictions. The optimization problem becomes:

**Figure 2:**
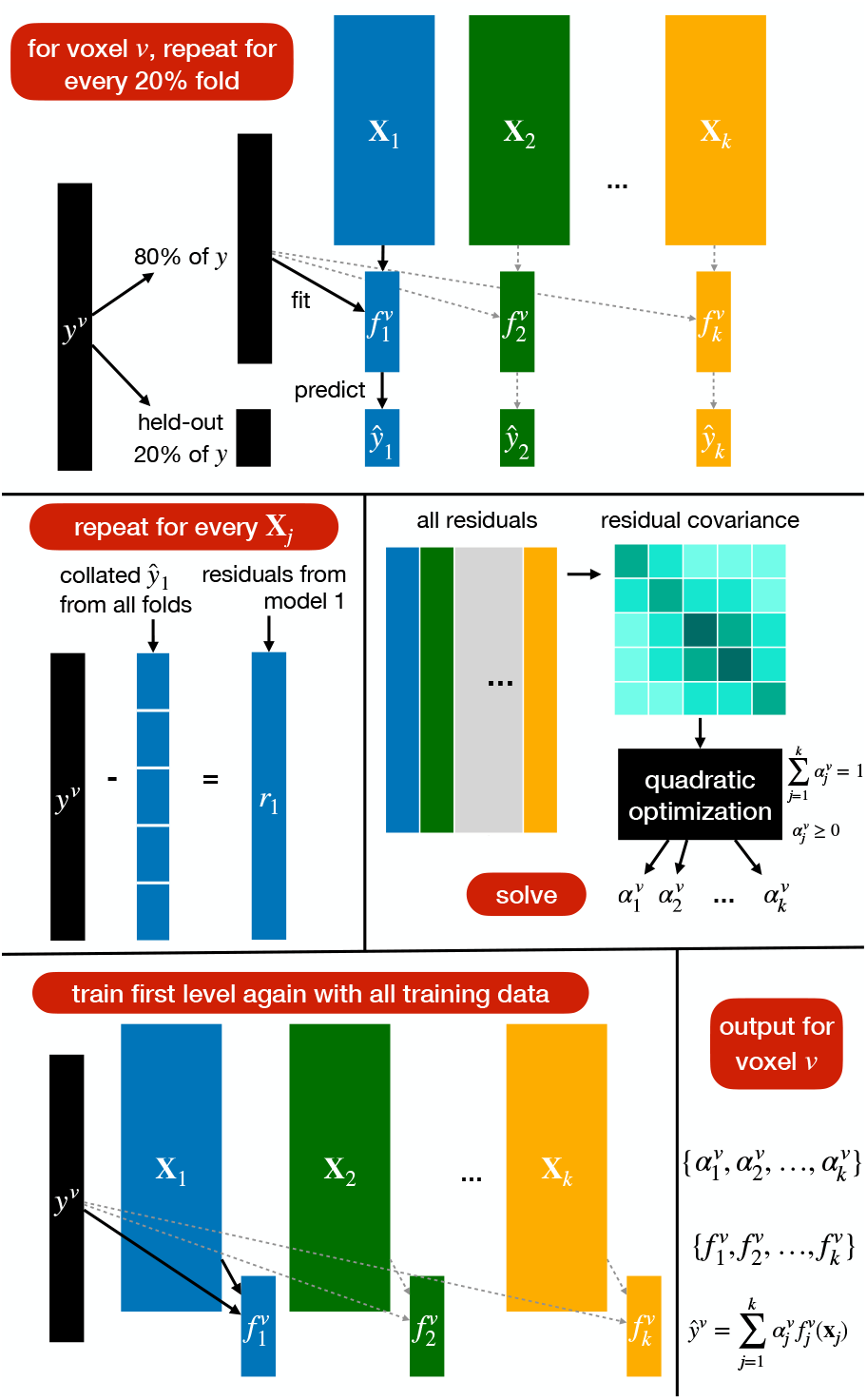
Schematic of the stacking method for each voxel *v*. First-level models are fit in cross-validation and used to produce out-of-set predictions for the entire training set. Residuals for each model are computed along with their covariance. The quadratic optimization problem is solved to obtain the 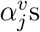. The 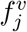 are fit again using the entire dataset.

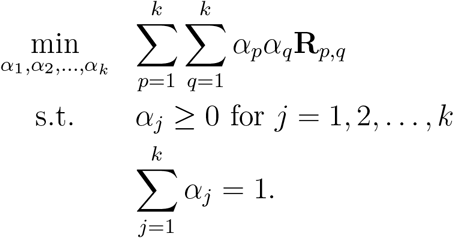

This minimization is set up to lead to high weights for more predictive feature spaces (small **R**_*i,i*_), while also selecting feature spaces that are not correlated in their prediction (small **R**_*i,j*_). Accurate feature spaces that have uncorrelated errors are complementary: for some samples, one will be more accurate than the other. By combining them, the aim is to improve prediction performance beyond their individual accuracy. For example, one might have two encoding models that yield the same absolute performance (e.g. *R*^2^ = 0.4) but actually capture distinct components of the variance in a voxel. Then, stacking can draw benefits by combining these models where where the error of each model might actually be compensated for by the other model. The constraints above are essential for the stacking procedure, and without them, the problem would be simplified to ordinary least squares (using the *ŷ*_*j*_’s as input), which might not have the same ensemble learning advantage that stacking brings [25]. Further, the summation to 1 makes the learned *α*_*i*_ easily interpretable into a weighted split of the prediction between the feature spaces.

We solve the above problem using the CVXOPT quadratic optimization package to obtain all *α*_*j*_’s [40]. If some amount of variance can be predicted from multiple predictors (owing to a shared component between the feature spaces), there exists an infinity of solutions to the optimization problem. However, CVXOPT is set up by default to break ties evenly between different predictors in case they contribute equally. This is advantageous for us, as it makes it so that our approach does not produce new inferences in favor of one of the feature spaces that are not justified by the data. We demonstrate in Supplementary Figure S1 a result where the inputs are two identical feature spaces (that is, they contribute to the final prediction equally). The weights are 0.5 everywhere, implying that all contributions are similarly weighted.

After solving the optimization problem, we estimate the *f*_*j*_s again using all the training data, and use the learned weights *α*_*j*_s to output 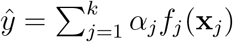. The entire procedure is repeated at each voxel. We summarize it in Algorithm 1 and in Figure 2.

#### Algorithm 1

Stacked encoding model training procedure

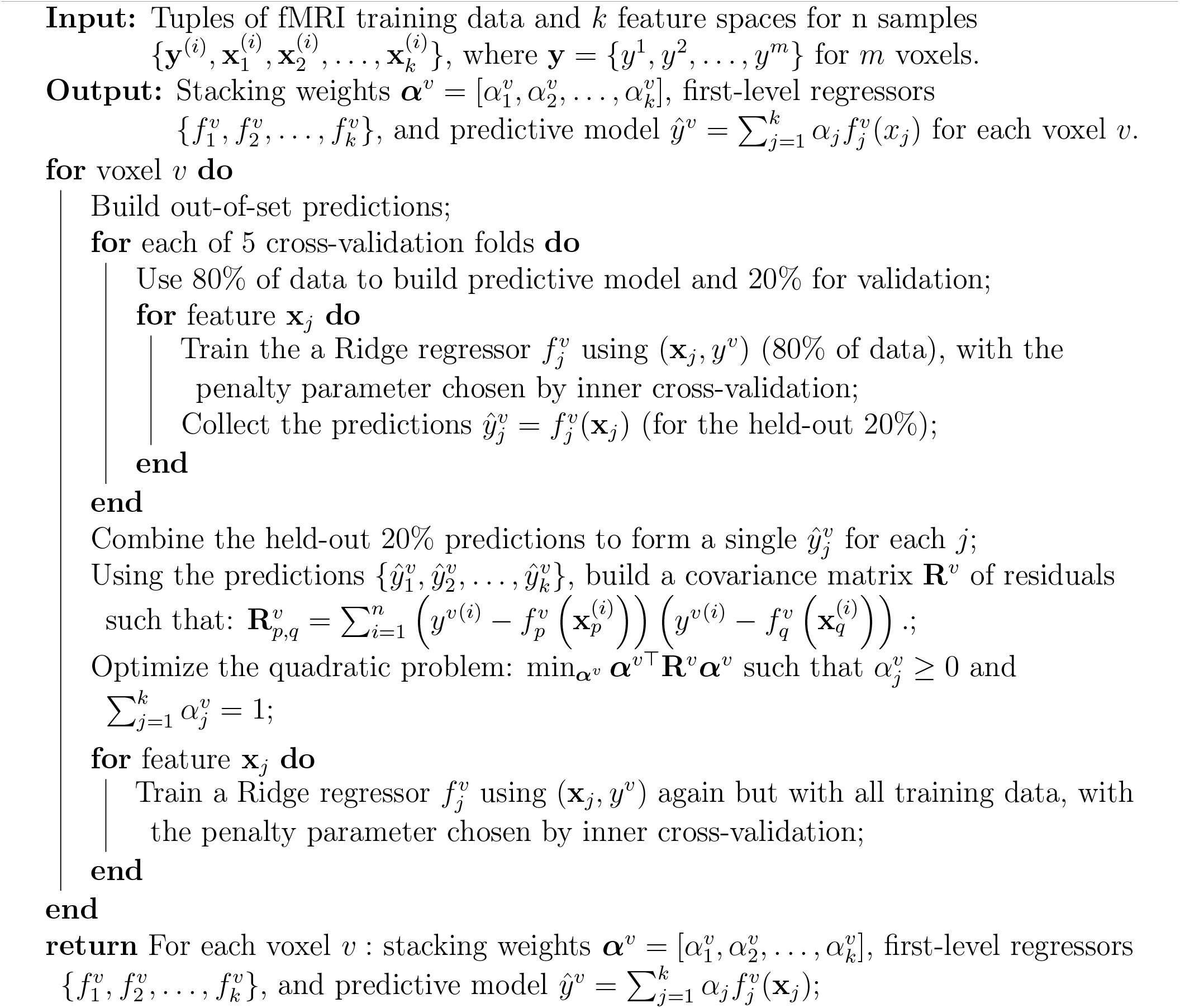

Note that in Algorithm 1, it is much more efficient to train the ridge regression estimators in parallel over all voxels. This is because, even though the parameter *λ*^*v*^ is estimated independently for each voxel, there are a lot of shared computations. Namely, for each of the potential tens of thousands of voxels, for each potential value of *λ*, the same (*X*^⊤^*X* + *λ*I) term should be inverted. (If another method is used to optimize the inversion, such as SVD decomposition, the computation is still repeated over all voxels.) Computing these once for each training set division and sharing the results over all voxels results in much faster computations, as is done in the Python package we share with this paper.

### Variance partitioning

Variance partitioning [34] is a popular approach to determine the relative importance of variables for regression models. Lescroart et al. [35] and de Heer et al. [36] apply variance partitioning to fMRI data. In addition to using estimates of prediction accuracy to compare feature spaces, the authors determined whether each feature space explains unique variance in the brain responses. To estimate the unique variance explained by a feature space, the variance explained when all feature spaces are used together (i.e. concatenated) is estimated. Encoding models that exclude each feature space are also built. Finally, set operations are used to capture the unique variance explained by each feature space.

Consider having two feature spaces **x**_1_ and **x**_2_, with each predicting variance *A*_1_ and *A*_2_ (respectively) of the data in one voxel. Using set operations, we find that the **unique** variance that **x**_1_ predicts is the following:

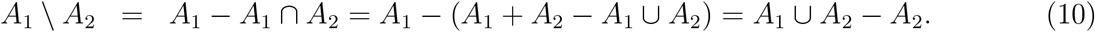

Here, *A*_1_ ∪ *A*_2_ is estimated using the coefficient of determination (*R*^2^) when evaluating the prediction performance of the joint model that uses **x**_1_ and **x**_2_ as inputs. *A*_2_ is estimated as the prediction performance (*R*^2^) of the model using only **x**_2_.

Similarly for multiple feature sets *{***x**_1_, …, **x**_*k*_*}*, the unique variance attributed to a single feature space **x**_*j*_ is:

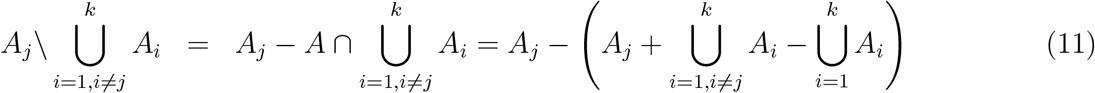

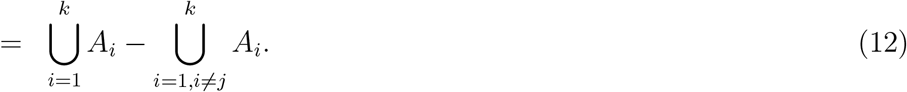

Variance partitioning can also be used for determining the unique variance explained by a union of feature spaces. Crucially, variance partitioning relies on accurately measuring performance when using any combination of feature spaces (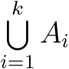 or the union of any subset of the *A*_*i*_s). For simplicity, let us consider variance partitioning with two feature spaces **x**_1_ and **x**_2_ as shown in equation 10. It is important that when evaluating the joint performance *A*_1_ ∪ *A*_2_, the measured performance is higher than or equal to each of the feature spaces (*A*_1_ and *A*_2_); otherwise the result will be nonintuitive (the union of feature spaces performs worse than an individual feature space, i.e., combining a feature space with another one reduces performance). This nonintuitive result also leads to a negative unique variance explained.

The typical approach to compute *A*_1_ ∪*A*_2_ has been to concatenate feature spaces. However, due to limitations in sample size with respect to feature space size, to confounds that might interact with the regularization or regression approach, or to noise, concatenating feature spaces might result in performance that is lower than the performance of a model with a subset of the feature spaces. In this paper, we show that using stacking to estimate the joint performance of multiple feature spaces is a more robust approach than simply concatenating the feature spaces. We show in simulation that stacking results most often result in joint model performance that is at least as good as the individual models. When using the high-quality fMRI data from NSD in conjunction with large feature spaces, we show that stacking results in better than or at least as good performance than the individual feature spaces, while concatenation often fails to do so. Indeed, one of our key points is that stacking is a better estimation method for variance partitioning because it more accurately estimates the performance of groups of feature spaces.

#### Structured Variance Partitioning

Our second contribution is structured variance partitioning. When more than two feature spaces are used, using variance partitioning can quickly become complex. For completeness, one might be compelled to compute the unique variance explained by all the subsets of feature spaces at disposal (all the *k* individual feature spaces, but also the 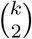 pairs of feature spaces, the 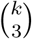 triples of feature spaces, etc.). This approach can quickly become intractable, and also often becomes too hard to interpret or lead to meaningful results.

Luckily, in many modern settings with many feature spaces, we do know a lot about the relationships between the feature spaces. Namely, in some settings where we use the different layers of the same network to model brain activity, we know exactly what the relationships between the layers are. For instance, in a usual convolutional neural network like AlexNet [33], *y*^*v*^ each layer **x**_*j*_ fully depends on the previous layer **x**_*j*−1_ and is obtained from it through a nonlinear function *g*_*j*_ that is learned when training the neural network. We want to determine how the information contained in those layers affects the brain response of different voxels. Recall that under specific assumptions that are satisfied in passive experiments such as image viewing (the experimenter manipulates the stimulus, and the stimulus precedes the response), the significant prediction performance of a brain region by an encoding model can be interpreted causally as indicating that some stimulus information contained in the feature space affects the activity in the brain region [41, 42]. From this observation, we are motivated to construct a more complex understanding of the effect of stimulus features on voxel activity (see Figure 3).

**Figure 3:**
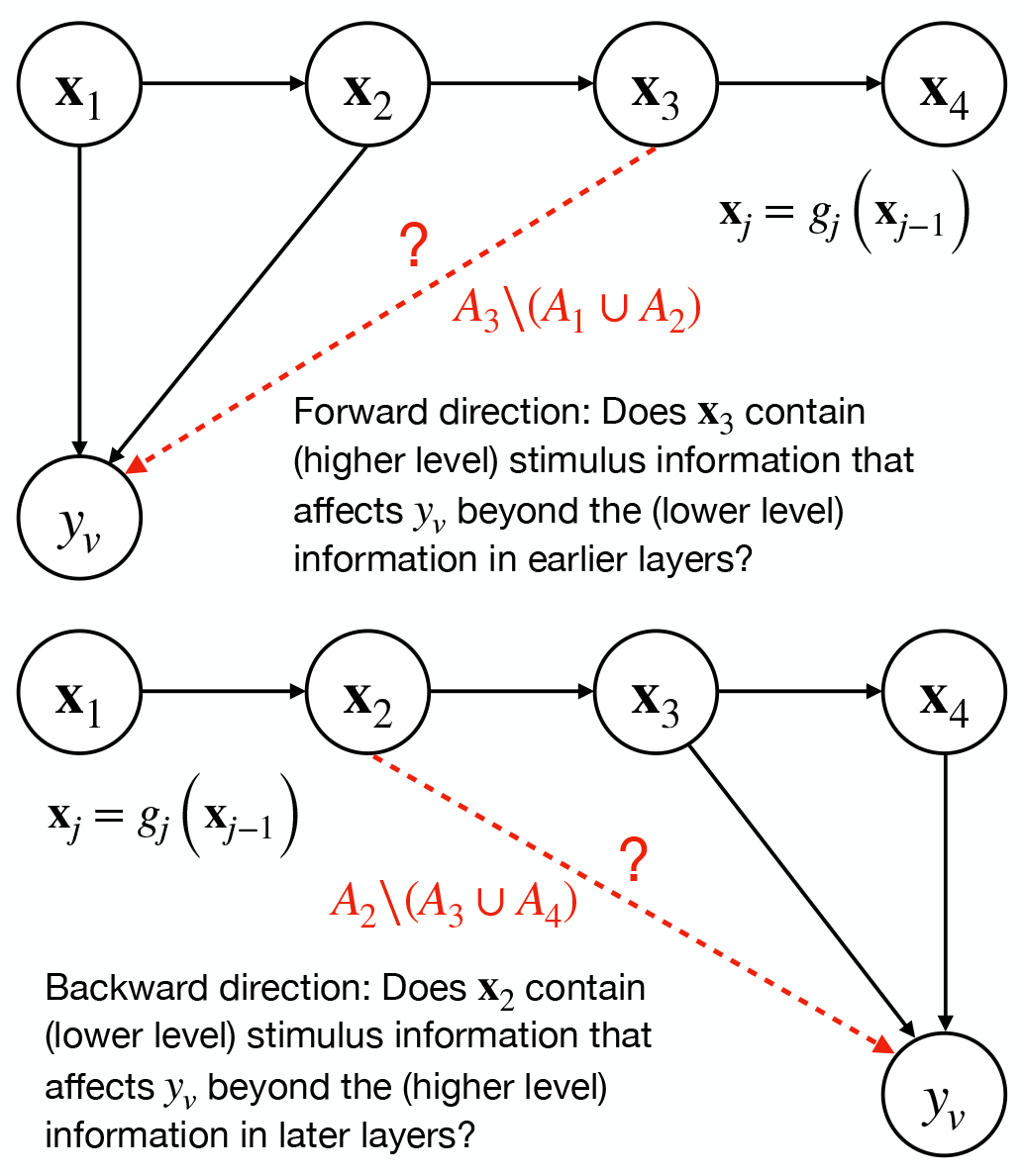
Illustration of two cases of structured variance partitioning: testing in the forward and backward direction.

We propose that knowledge about the structure of feature spaces relationship can be embedded in our inference pipeline. Knowing that the layers are functions of each other, we can refine the variance partitioning analysis. For example, looking at the unique variance for each layer might be misleading because of the large amount of shared information between layers given that they are functions of each other, and thus the unique variance for each layer at a voxel would look small for all layers, and the result would be hard to interpret. However, we can ask a more structured question, such as: How complex (i.e., high-level) do the representations need to be to predict a specific voxel? Specifically, we can ask, for a hypothetical 4-layer network, do we need to go up to layer 3 to predict voxel *v* (see Figure 3[Top])? In other words, given layer 1 and 2, is there still higher-level information contained only in layer 3 (and not in layers 1 and 2) that can predict unique variance in voxel *v*? In terms of interpretation, this is akin to asking if voxel *v* is modulated by stimulus features at least as complex as ones contained in layer 3.

To answer this question, we construct a series of “forward” variance partitioning analyses, where we compute the unique variance explained by every layer *j* (2 ≤ *j* ≤ *k*) beyond the lower layers, i.e. we compute 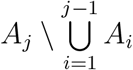. Again, the performance of any group of features 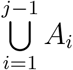 is estimated as the prediction performance (*R*^2^) of the joint models that uses **x**_1_ … **x**_*j*−1_. This returns *k* − 1 results for each voxel, allowing us to determine whether each layer above layer 1 can predict unique variance on top of the earlier layers. To combine this information into a single statistic that allows us to visualize information on the brain and interpret it, we look at the series of stacked performances 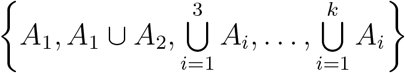, which are computed as a biproduct of performing all the variance partitioning analyses 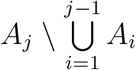. Recall that because we are combining the sets using stacking, we expect that this series would be increasing, with every element being at least as large as the one before. Thus, the very last element 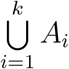 should correspond to the highest prediction accuracy that can be achieved for voxel *v* using feature spaces *{***x**_1_, …, **x**_*k*_*}*. We ask, for voxel *v*, what is the latest layer *j* we need to include so that we reach at least 95% of the top performance? I.e., we find:

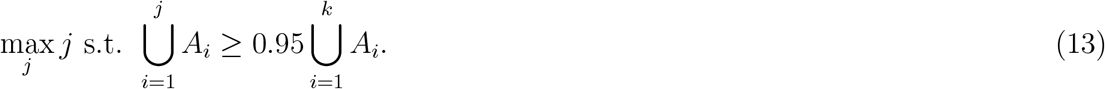

We may also be interested in a separate question. When looking at a high-level area which we know processes high-level information, we might still be interested in knowing if it also processes low-level information, and if so to what level (see Figure 3[Bottom]). To answer this question, we construct a series of “backward” variance partitioning analyses, where we compute the unique variance explained by every layer *j* (1 ≤ *j* ≤ *k* − 1) beyond higher layers, i.e., we compute 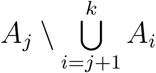 (again using *R*^2^ to assess prediction performance for any group 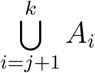). To compute a single statistic per voxel, we consider the series of stacked performances 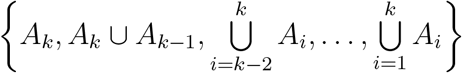. We also expect that this series would be increasing, with the last element corresponding to the top prediction accuracy. We ask, for voxel *v*, what is the earliest layer *j* we need to include so that we reach at least 95% of the top performance? I.e. we find:

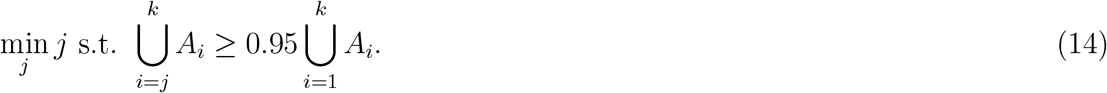

By using both the forward and the backward directions, we can construct an interval of the features that are processed at a voxel, with the backward direction providing a lower bound on the layers that are relevant for the voxel, and the forward direction providing an upper bound. This interval offers more information than the simple choice of a top layer per voxel. It has been observed that the different layers of the same model overlap in their ability to predict [18]. Furthermore, in the domain of human vision, it has recently been shown that different parts of the early visual visual cortex have complex organization and do not necessarily process information serially in direct correspondence with the layers of a convolutional network [43], and thus, our structured variance partitioning can serve as a better tool for performing network alignment. We have used an ordered organization to perform structured variance partitioning; however, different setups and other relationships between feature spaces might call for a different organization.

### Evaluation

We evaluate the stacking approach by comparing it with the baseline of concatenating the feature spaces into a larger feature space. In our simulated experiments, we compare the performance of both estimation approaches in predicting responses (using *R*^2^), as well as the error of each model in variance partitioning (in the simulated experiment, the true unique variance is known).

Using real data, we compare the prediction performance of the stacked models to the concatenated models, and to the best performing out of the individual models (which use only one feature space as input). This last comparison evaluates whether stacking and concatenations are appropriate for variance partitioning, as the joint model (either stacked or concatenated) should perform at least as well as the best individual model.

We also use the stacked and concatenated models to perform structured variance partitioning (combining the forward and backward directions). We evaluate the constructed maps in terms of their smoothness and coherence, and the hypotheses they suggest about the brain. Throughout this paper, we evaluate the stacking approach by training and testing on the entire dataset (in simulation or with real data) in an outer cross-validation setting. At each fold, we hold out 20% of the dataset and use 80% as training data (input to Algorithm 1). The results are combined over all the folds. We evaluate the concatenated model in the same fashion (80% of the data is used to train a Ridge regressor with the penatly chosen by inner cross-validation at each voxel). The results are combined over all folds.

### Simulated experiments

To evaluate the performance of our stacking approach, we design a set of simulations in which we control different parameters that can be relevant for fMRI experiments. The steps to simulate data are as follows:

- For each feature space *j*, we sample a feature space matrix 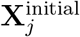 with *n* samples and dimension *d*_*j*_. Each row of **X**^initial^ is from a multivariate Gaussian with zero mean and a Toeplitz covariance matrix **∑**_*j*_, defined using an exponentially decaying function (**∑**_*j*_ (*u,v*) = exp(−(*u* − *v*)^2^*/*(*s × d*_*j*_))), where *s* is a scaling factor). This allows for some simple dependence structure between the features in a feature space.
- 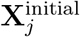 and 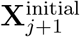 are independently sampled. For *j* ∈ *{*1, 2, …, *k* − 1*}*, we further create a correlated component that is shared between each two consecutive feature spaces. We sample the matrix 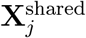 with *n* samples and dimensions *d*_*j*_ (also using **∑**_*j*_). We further sample a rotation matrix **O**_*j*_ of size *d*_*j*_ *× d*_*j*+1_ (each column is sampled from **∑**_*j*_). We create the feature spaces 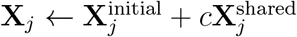 and 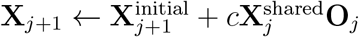, where *c* is a chosen factor that governs the correlation between the feature spaces.
- We sample **w**, a long weight vector corresponding to all features, from a multivariate normal with mean 0_*d*_ and a Toeplitz covariance matrix **∑**_*d*_ where 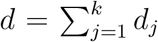. We divide **w** into the individual feature spaces (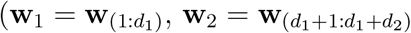, etc.).
- We set by hand the *α*_*j*_, the contribution of each feature space under the stacking model 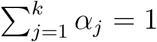 and *α*_*j*_ ≥ 0, *j* = *{*1, …, *k}*).
- We sample the noise component ***ϵ***, a vector of *n* numbers, each sampled from a normal distribution with mean 0 and variance 1. We set *σ* as the scaling factor for the noise term (equivalent to sampling from a normal distribution with mean 0 and variance *σ*^2^).
- Finally, we compute **y**, the vector of simulated voxel activity as:

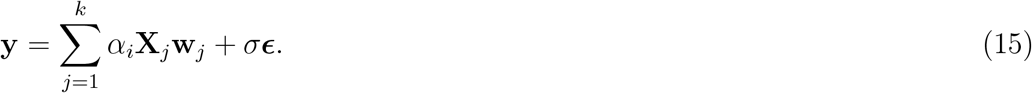

We simulate an experiment with 4 feature spaces *{***X**_1_, **X**_2_, **X**_3_, **X**_4_*}* with each pair **X**_*j*_, **X**_*j*+1_ having a correlated component (*c* = 0.2). For evaluation, we compare the prediction performance (*R*^2^) of stacked models to the performance of concatenated models, and to the maximum individual performance of individual models using each of the feature spaces. We also judge the models on the basis of their ability to estimate the unique variance explained by **X**_1_, i.e., 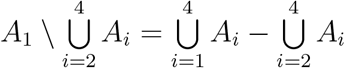. We compute 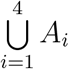 and 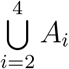 using either stacking or concatenation. Because we are simulating the variables, we have access to the initial component of **X**_1_ that is not correlated with **X**_2_, and thus we can use it to obtain the true value of the unique variance explained by **X**_1_, by computing 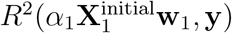. We compute the absolute error between this value and its estimate from the stacked and concatenated models. We vary one of the four parameters in Table 1 at once. For each parameter setting, we use 50 samples. We include our simulation code in our code package.

**Table 1:** Parameters controlling data simulation

| Parameter | Definition |
| --- | --- |
| $d_j$ | dimension of $\mathbf{X}_j$ |
| $n$ | number of samples |
| $\sigma$ | noise level |
| $\alpha_j$ | Stacked weights |

By varying *n*, the number of samples, with respect to the dimensions of the feature space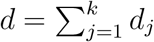, we can assess the complexity of the learning problem and the relative number of data points that are required to obtain robust performance. By varying the *α*_*j*_’s, we can explore the different settings of relationships between feature spaces and a voxel, and the behavior of our approach under each. By varying *d*_*j*_ in conjunction with *α*_*j*_, we can test whether important feature spaces with few feature dimensions could still be detected when compared with larger feature spaces with less importance. By varying *σ* we can study the robustness of our method under noise. The relevant parameters in our simulation are summarized in Table 1.

### The Natural Scenes Dataset

The Natural Scenes Dataset (NSD) by Allen et al. [1] is a large-scale fMRI dataset conducted at ultra-high-field (7T) strength which consists of whole-brain, high-resolution (1.8-mm isotropic, 1.6-s sampling rate) fMRI measurements of 8 healthy adult subjects. It is shared with a nonexclusive, royalty-free license for research and educational purposes. While fMRI responses are recorded, subjects view thousands of natural color scenes over the course of 30-40 scan sessions. While viewing these images, subjects are engaged in a continuous recognition task in which they fixate at the center of the screen and report whether they have seen each given image at any point in the experiment. Each image is repeated up to three times and the average of the repeats is calculated. A general linear model was used to estimate single-trial beta weights. We used the betas_fithrf_GLMdenoise_RR preparation of the betas. Using FreeSurfer [44, 45], cortical surface reconstructions were generated to which the beta weights were mapped. One averaged fMRI response to each image was obtained by z-scoring the beta weights per voxel for each run and then averaging across repetitions of the image. All brain visualizations were produced using the Pycortex software [46].

### Visual feature extraction

Agrawal et al. [47], Eickenberg et al. [48], Tripp [49], Zhang et al. [50], Pasupathy et al. [51], Mohsenzadeh et al. [52], and others show that different layers of AlexNet [33] can predict a hierarchical set of regions in the visual cortex. We replicate this analysis and extract representations from the five convolutional layers and two fully connected layers of AlexNet for all the stimulus images. Due to memory limits, we preprocess the features we extracted using PCA before the analysis. We reduce the dimensions of all feature spaces to 1024. Supplementary Figure S2 shows the similarity across images for each pair of feature spaces.

### Computational Resource

All experiments are carried out on a workstation with an Intel Xeon W-2145 CPU @ 3.70GHz and one NVIDIA 1080-Ti GPU. Image feature extraction for the NSD dataset is conducted on the GPU. Training of the stacked encoding models is conducted on the CPU.

## Results

### Simulated data: Stacking appears advantageous in multiple settings

In this section, we evaluate stacking on simulated data by varying four parameters that can govern the statistical properties of an encoding model analysis. Feature spaces with small number of features but that are important for a voxel might be more difficult to identify. We first vary the dimensionality of one of the feature spaces (**X**_1_) while keeping the dimensionality of the other feature spaces constant (Figure 4(a)). This allows us to test whether stacking detects important feature spaces more reliably than concatenation even when they have small dimensions. Here, we know the ground truth because we have simulated the data, thus we know exactly what is the unique variance explained by **X**_1_. We can thus directly compare the performance of the stacking approach to the more classical approach of concatenating feature spaces. Under our settings, we find that, when estimated during stacking, the unique variance explained by **X**_1_ has much less error than when estimated using concatenation. For some settings, the error is minimal for stacking and goes beyond 50% of the variance for concatenation (see Figure 4(a), right most panel). The advantage of stacking over concatenation gradually reduces when the dimensionality of **X**_1_ is increased.

**Figure 4:**
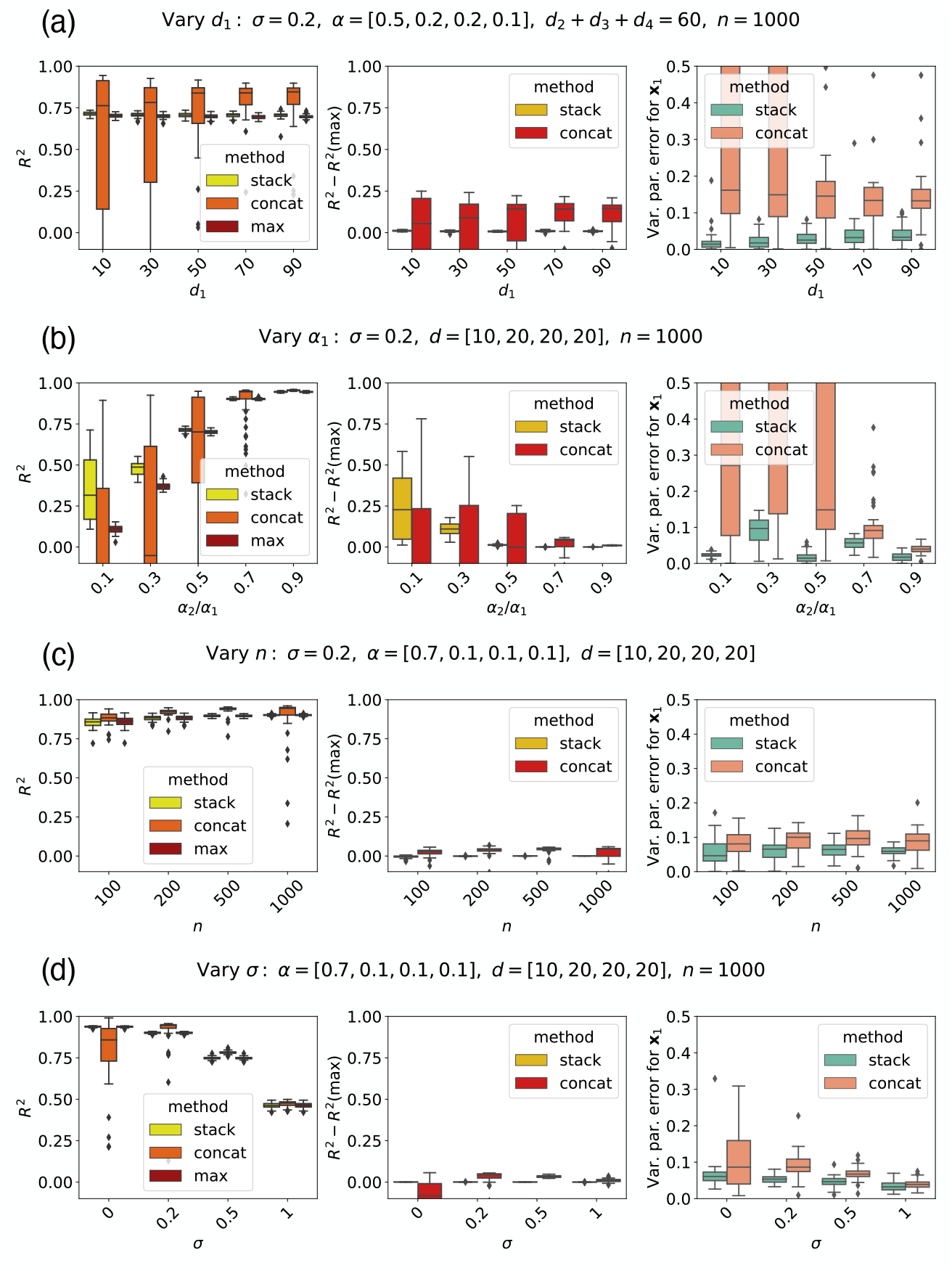
Simulation results. **(a)** Varying *d*_1_ while keeping *{d*_2_, *d*_3_, *d*_4_*}* fixed. The stacked model has much less error than the concatenated model when estimating the unique variance explained by **X**_1_. **(b)** Varying *α*_1_. The stacked model has much less error than the concatenated model when estimating the unique variance explained by **X**_1_, and the advantage reduces with increasing *α*_1_. **(c)** Varying *n* (*α*_1_ = 0.5). The stacked model maintains modest improvement in error in unique variance explained by **X**_1_, while the concatenated model has a modest advantage in *R*^2^. **(d)** Varying *σ*. The stacked model shows an advantage over the concatenated model, most pronounced for small *σ*.

Feature spaces with an intermediate importance might also be difficult to identify in the presence of other feature spaces. Next, we vary *α*_1_, the importance of **X**_1_ during the generation of activity (Figure 4(b)). This allows us to test whether stacking correctly estimates the important feature spaces even when they don’t have a very high importance, and other feature spaces contribute to most of the activity. Under our settings, we find that, when estimated during stacking and not concatenation, the unique variance explained by **X**_1_ has much less error, and that this advantage is reduced the higher the importance of **X**_1_. For very high importance for **X**_1_, the performance of the concatenated model becomes indistinguishable from stacking.

Having a small dataset might interfere with our ability to identify the importance of feature spaces. We vary *n*, the number of samples available (Figure 4(c)). This allows us to test whether stacking has an advantage over concatenation for small sample sizes. Under our settings, we find that for low sample sizes, the stacked model has some small improvements over the concatenated model in estimating the unique variance of **X**_1_, while the concatenated model has higher prediction performance (*R*^2^). Even in cases where the stacked model performance is not as high as the concatenated model performance, it is still higher or very close to the maximum model performance.

Finally, noise might interfere with our ability to identify the importance of feature spaces. we vary *σ*, the noise level (Figure 4(d)). This allows us to test whether stacking has an advantage over concatenation in the presence of noise. Under our settings, we find that stacking does have an advantage over concatenation in estimating the unique variance of **X**_1_, but that this advantage is most pronounced in low- to mid-level noise, and is greatly reduced for high-level noise.

In summary, in the specific conditions we simulated, the advantage of stacking over concatenation in estimating the unique variance explained by **X**_1_ is greatest when **X**_1_ has a small number of dimensions (Figure 4(a)), and when *α*_1_ ≤ 0.5 (Figure 4(b)). This could be due to the second-level optimization procedure used in stacking, which takes only a small number of inputs (one prediction from each feature space vs. a large number of features from each feature space) and thus might better able at identifying the importance of a feature space with small dimensions or relatively small importance. The advantage of stacking persists across different sample sizes (Figure 4(c)), but reduces with added noise (Figure 4(d)). As we mentioned above, for some simulation settings, the concatenated model performance is reliably greater than the stacking performance; however, for those settings, the concatenated model has a greater error in estimating the unique variance explained by **X**_1_.

Another important observation is that, under our settings, the performance of the stacked model is reliably similar to or higher than the performance of the maximum individual model. Further, for some simulation settings (e.g. low dimensionality of **X**_1_ and low importance *α*_1_), the concatenated model performance sometimes is much larger than the maximum model performance, but for the same simulation settings, it can also be much lower than the maximum model performance. As explained previously, for variance partitioning, it is best to use a method that produces results when combining feature spaces as least as good as the performance of individual the feature spaces. Thus, stacking is suggested by those results to be more useful for variance partitioning.

The simulation experiments we show in this section are a small sample of the infinity of simulation experiments that can be produced. We use these few experiments to begin to understand the behavior of stacking with respect to concatenation, but they are not a proof or a guarantee that stacking will always be more accurate. If the experimenter is working in a regime where it is not clear that stacking has an advantage and they are only interested in improving prediction accuracy, one solution is to add an additional feature space **X**_*k*+1_ = [**X**_1_, **X**_2_, …, **X**_*k*_], which is the concatenation of the individual feature spaces, and use stacking with *{***X**_1_, …, **X**_*k*+1_*}*. In the case where the concatenation has an advantage, it should be conferred to the stacking model: in that case the model should mostly select the concatenated feature space. This selection would render the individual feature spaces redundant, but would improve prediction performance. Variance partitioning can still be performed in this case (a concatenated feature space is added to each stacked model by concatenating only the feature spaces in that stacked model).

### NSD: Stacking improves encoding model performance

The first question we address here is: does our stacking method improve the encoding model performance with real data? Again, we use the coefficient of determination (R^2^) for evaluation of the ability to predict fMRI data per voxel. We compare the performance using our stacking method to the performance of an encoding model that uses concatenation to combine feature spaces. We compare the performance of both approaches to the maximum encoding performance of ridge regression models that each use one of the individual feature spaces (i.e., the first-level models in stacking). We compute R^2^ on held-out data in a 5-fold cross-validated way (we run the entire stacking approach in Algorithm 1 in a nested manner, and predict the held-out data).

Figure 5 shows our encoding result for subject S1 in the NSD dataset (Supplementary Figures S3-S10 show the results for all subjects). We find that the stacked model performance is very high in the primary and high-level visual cortex, along with some scattered regions in the prefrontal cortex. The stacking performance is higher than the concatenated model performance (Figure 5(e)) and the maximum individual feature space performance (Figure 5(d)), which also follow a similar pattern of being very high in the primary visual cortex and high in the high-level visual cortex (Figure 5(b) and (c)). Importantly, across the top 2000 voxels (shown in Figure 5(g)), the stacked model performance is always higher than the maximum individual model performance (and for the rest of the voxels, the stacked performance is also higher or equal to the maximum performance, up to rounding error), whereas the concatenated model performance is very often lower than the maximum individual model performance. As highlighted previously, this suggests that the stacked model is a more useful method for variance partitioning, since it results in performance that is greater than the individual encoding model performances.

**Figure 5:**
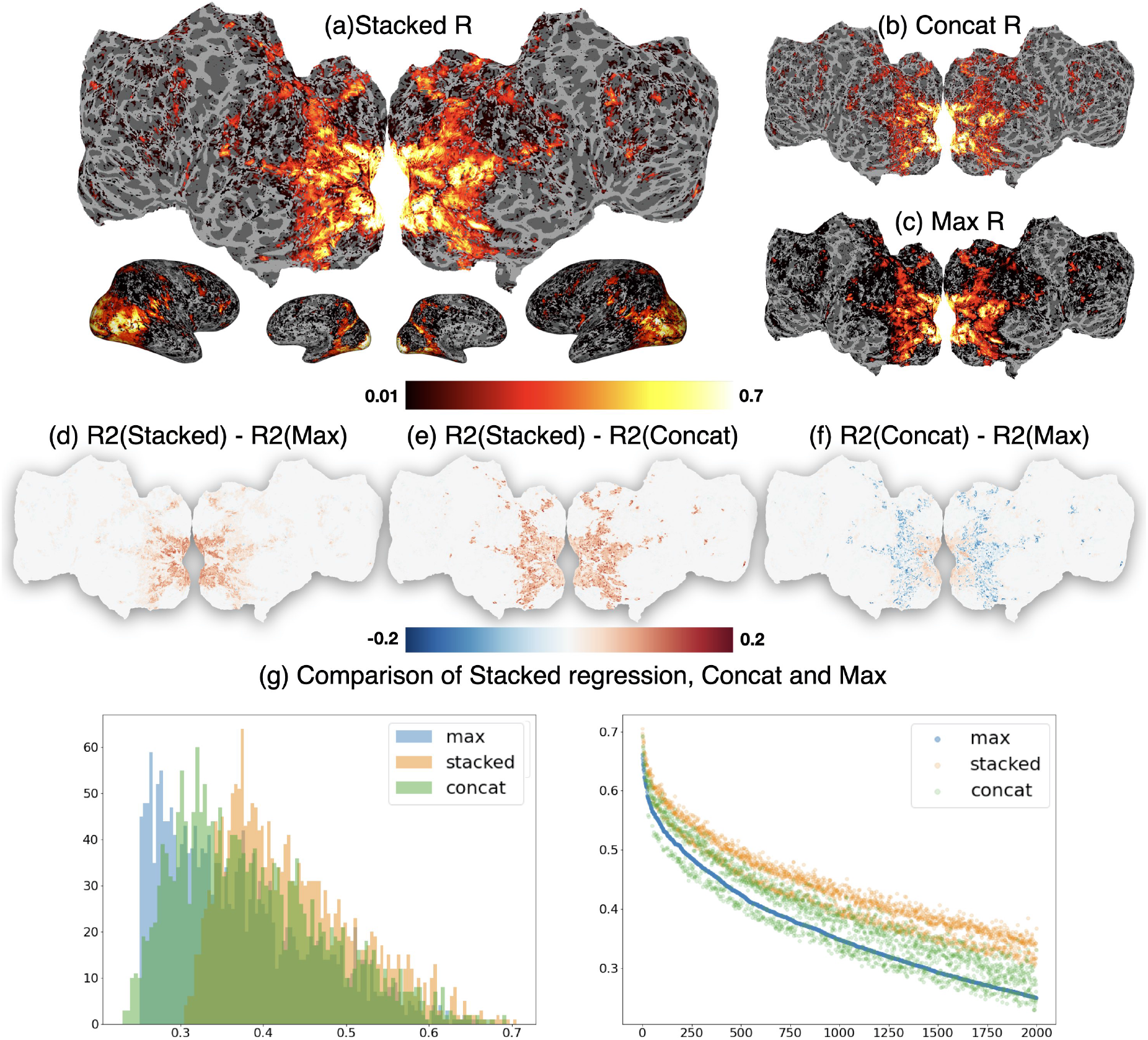
Encoding performance for subject S1 in the NSD dataset. **(a):** Prediction performance is evaluated using R (The square root of *R*^2^, the coefficient of determination, when *R*^2^ *>* 0) for the stacked encoder using all layers of AlexNet to predict the brain activity related to viewing natural scenes. We show the prediction performance on the flattened brain surface (top) and the inflated brain surface (bottom, with the views in order as right lateral, right medial, left medial, and left lateral). **(b)** R for the encoding model that uses as input feature space the simple concatenation of all the layers of AlexNet. **(c)** Maximum R of the encoding models that use as input a feature space an individual layer of AlexNet. Panels **(d**,**e**,**f)** show the difference between *R* of the stacked model and the maximum individual *R, R* of the stacked model and *R* of the concatenated model, and *R* of the concatenated model and the maximum individual *R*, respectively. For all three *R* values plotted in **(a**,**b**,**c)**, the primary visual cortex is remarkably well predicted with values close to 0.7. The high-level visual regions are also well predicted. However, the *R* of the stacked model is higher than the *R* of the concatenated model and the maximum individual *R* in all well predicted voxels. However, the *R* of the concatenated model is higher than the maximum individual *R* in the primary visual cortex, but lower than it in the high-level visual regions. This can also be seen in **(g)** through a histogram (left) and a plot (right) of the three values of the top 2000 performing voxels, ordered by their decreasing maximum individual *R* value.

### NSD: Stacking generates interpretable brain maps

We have shown that our method improves prediction performance. Nevertheless, in science, the main goal is to provide interpretability. Next, we show that we can generate more interpretable brain maps using our stacking method.

We build a feature attribution map by assigning to each voxel the feature space that predicts it best. For the stacking results, we pick the feature space with the highest *α*_*j*_. We compare this feature attribution method to choosing the best feature space for a voxel as the feature space that results in the best performance when used on its own in an encoding model.

Figure 6 shows our feature attribution result for S1, S2 and S3 (Supplementary Figures S3S10 show the results for all subjects). We have seven features extracted from AlexNet for each subject, as we mentioned above.

**Figure 6:**
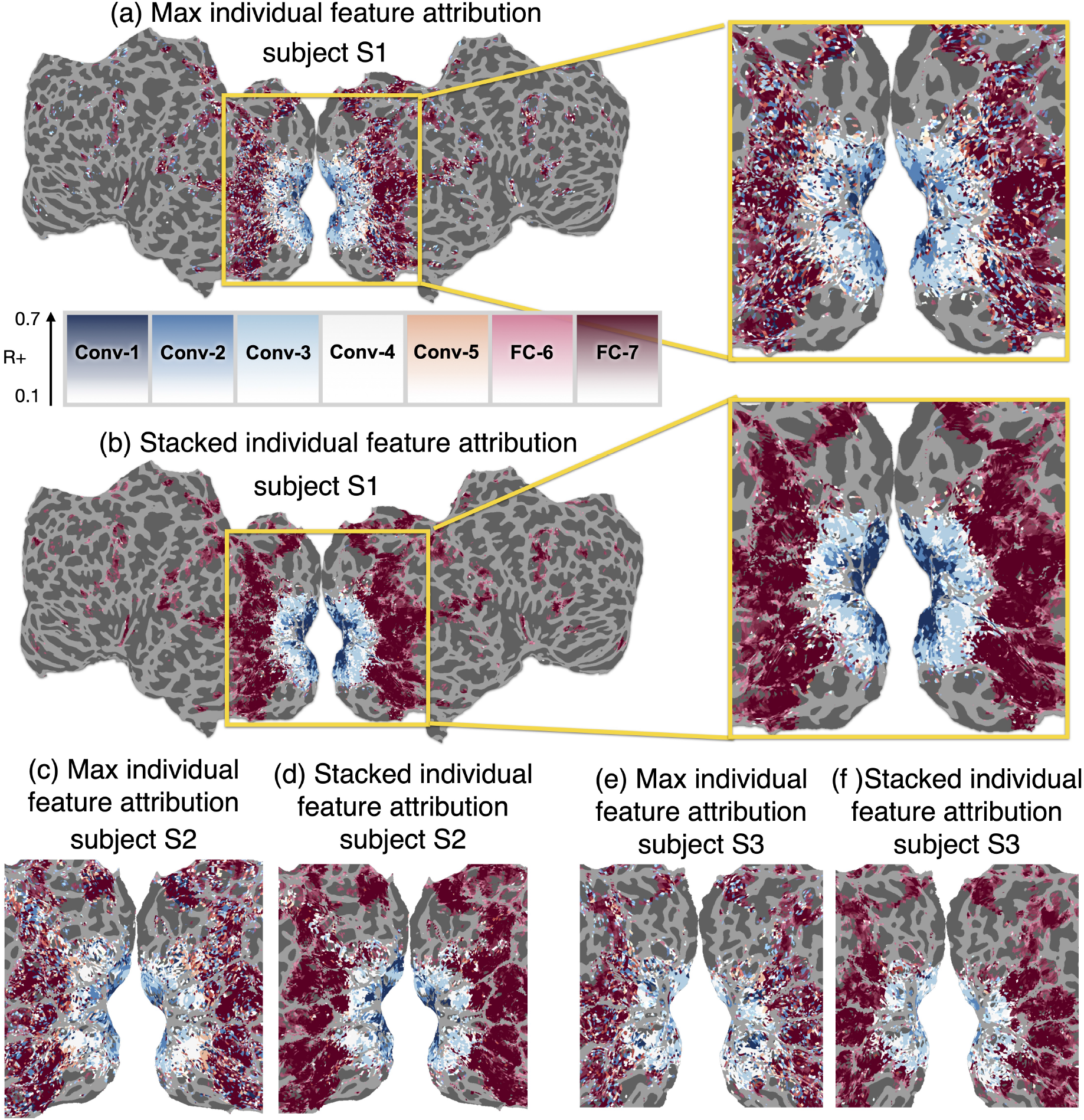
Feature attribution results in the NSD dataset. For each voxel, we pick the “best” feature space using two approaches. **(a)** We pick the best feature space as the feature space with maximal encoding performance *R* at a voxel. The result is shown on the flattened brain of subject S1, and a zoom-in plot shows the primary and high-level visual cortices. The color corresponds to the layer of AlexNet and the transparency level corresponds to performance, with high *R* corresponding to completely opaque colors. **(b)** For subject S1 again and the same visualization parameters as **(a)**, we pick the best feature space as the feature space that has the highest *α*_*j*_ in the stacking model of a voxel. Both results in **(a)** and **(b)** show a correspondence between the layers of AlexNet and the hierarchy of visual regions, as reported by Agrawal et al. [47], Güçlü and van Gerven [17], Eickenberg et al. [48], and others. However, the feature attribution using the stacking method results in much smoother maps, even though each voxel is optimized separately and no spatial smoothness is enforced. Spatially smooth results are more biologically plausible given what we expect about the hierarchy of the visual system. We see similar results for subject S2 **(c**,**d)** and subject S3 **(e**,**f)**.

Consistent with previous results, we find that the successive layers of AlexNet predict the areas in the primary visual cortex following the hierarchy of the visual system [48, 53, 54, 17, 55, 56, 57, 19, 58]. The early convolution layers in AlexNet are chosen for the early visual cortex (including V1, V2); meanwhile, top fully connected layers in AlexNet are more likely to be chosen for high-level regions, including ROIs selective for places or faces. While both the maximum feature space attribution and the stacked weight attribution show a similar result, the stacked attribution follows a much smoother and spatially consistent pattern, while the maximum feature space attribution is grainy (e.g., some sparse voxels in high-level visual cortex are assigned to very early layers such as Conv-1 and Conv-2). Given the expectation that the fMRI results would be smooth in space, the stacked results appear to be more robust and more reliable. Recall that the stacked approach does not enforce spatial smoothness and that the observed smoothness is purely a function of the improved fitting method.

### NSD: Structured variance partitioning allows detailed hypothesis testing

Looking at the maximal stacking weight per voxel led to smooth maps; however, it is not always the most informative metric. As discussed above, there might be a strong relationship between feature spaces, and a more informative approach is to account for these relationships. We show in this section the results of structured variance partitioning using the forward and backward direction, and using these to establish an interval for each voxel that captures the range of visual features (as operationalized through AlexNet) this voxel is sensitive to.

We perform structured variance partitioning in the backward and forward directions introduced in the methods. Specifically, in the backward direction, we start at the deepest layer of AlexNet (FC-7) and determine for each voxel, how many earlier layers need to be added to achieve at least 95% of accuracy achieved with all layers (Figure 7[Top]). In the forward direction, we start at the shallowest layer (Conv-1) and similarly determine how many subsequent layers need to be added before reaching at least 95% of accuracy achieved with all layers (Figure 7[Top]). This creates an interval of layers that appear necessary to predict a voxel’s activity.

**Figure 7:**
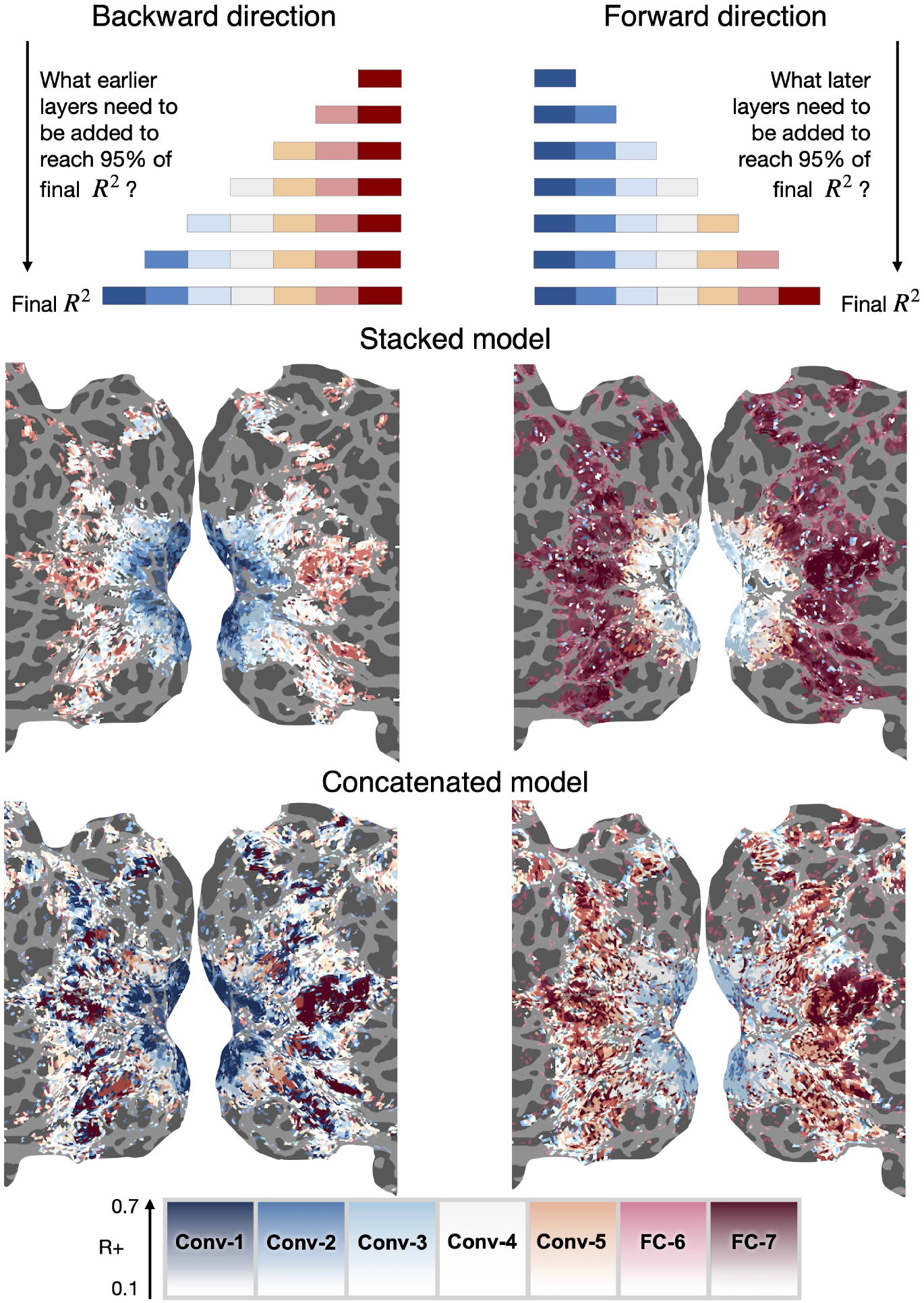
Structured variance partitioning in the NSD dataset. In the backward direction, we determine for each voxel, starting at FC-7, how many earlier layers we do have to add before achieving at least 95% of the accuracy achieved with all layers. In the forward direction, we start at Conv-1 and similarly determine how many later layers need to be added. Using the stacked model, the results in both the backward and forward directions are spatially smooth and show a logical interval (the backward layer is earlier than the forward layer). For voxels in the earliest part of the visual cortex (corresponding to V1), layers Conv-1 to Conv-3 appear to be required for prediction. For regions in high level visual areas such as the EBA (see Supplementary Figure S11 for atlas), layers FC-6 and FC-7 appear necessary, while in other parts of high level visual cortex, such as the PPA, layers Conv-4 to FC-7 appear to be required. Results using the concatenated model are much less spatially smooth. The intervals of prediction are either very broad or too narrow (some parts of high level visual cortex appear to need all layers while other ones right adjacent to them are mostly explained by Conv-4).

Using the stacked model, the results in both the backward and forward directions are spatially smooth (Figure 7[Middle]). The layers chosen by the backward and forward procedures show a logical interval (the backward direction layer is earlier than the forward direction layer). For voxels in the earliest part of visual cortex (corresponding to V1, see the Supplementary Figure S11 for the atlas), the layers Conv-1 to Conv-3 appear to be required for prediction. For regions in high-level visual areas such as the extrastriate body area (EBA, Supplementary Figure S11), layers FC-6 and FC-7 appear to be required for prediction, while in other parts of high-level visual cortex, such as the parahippocampal place area (PPA, Supplementary Figure S11), layers Conv-4 to FC-7 appear to be required. The results using the concatenated model are less spatially smooth. While the broad pattern of results obtained with the concatenated model resembles stacking, the resulting intervals are either very broad or too narrow (some parts of high level visual cortex appear to need all layers while other ones right adjacent to them are mostly explained by Conv-4). These results can also be analyzed by plotting the interval width of the layers required to predict a voxel (the result of the forward direction minus the result of the backward direction), and can be seen in the Supplementary Figure S12. This figure shows that the concatenated model leads to many nonintuitive results (regions where the interval is negative). This figure also shows that, using the stacked model, some high-level ROIs such as EBA and FFA, as well as some regions in early visual cortex, have a small interval width (only one or two layers are required to predict them), while intermediate regions of the visual system and high-level place regions (RSC, OPA and PPA) require up to four and five layers. The fact that the high-level visual cortex is known to be involved in high level semantic processing [59, 60, 61, 62, 63, 64, 65, 66, 67], and that the place regions specifically are also tuned for intermediate level features [35, 68, 69], support the conclusions from the stacking results.

## Discussion

The first contribution of this paper is to propose using stacked regressions to predict brain activity from multiple feature spaces. The stacking model learns the parameters of a convex combination at each voxel, that can be interpreted as the importance of each feature space for predicting that voxel. We showed in simulation that stacking can be more accurate than simply concatenating the feature spaces in multiple settings. We also show that when using the NSD data and feature spaces from AlexNet, stacking leads to more robust results. This suggests that stacking might be useful in general for modeling large fMRI datasets of good quality with high-dimensional feature spaces from neural networks. However, as with every statistical method, stacking is not a one-size-fits-all tool, and the decision to use it should be made with consideration of the problem at hand and critical thinking.

We have implemented our first-level encoding models that take one feature space as input as linear predictors. However, the stacking method, can be used with nonlinear predictors of arbitrary complexity (including, for example, encoding models that are made of deep neural networks instead of linear predictors). As we have implemented them, our stacked models are effectively linear models that are regularized under a specific set of constraints, which we show can be advantageous for building encoding models of fMRI data.

The stacking weights we learned while training our method provide us with a consistent and smooth feature attribution without any spatial constraint. These smooth feature attribution results make biological sense since the brain’s organization is thought to be smooth. We argue, however, that a more informative interpretation method than looking at the top feature space per voxel—which presents an incomplete picture and can be more prone to noise—is to perform structured variance partitioning.

Structured variance partitioning is the second method we proposed. It allows a more complex understanding of a voxel by considering its relationship to multiple feature spaces, while incorporating the relationship between these feature spaces. We think structured variance partitioning is a useful tool to deal with the complexity of modern encoding models, which often rely on feature spaces from large neural networks, and which are often used to model complex data acquired in rich, naturalistic experiments. The layers of a deep neural network are functions of each other, thus they can be quite correlated. Consequently, they might map onto the brain in a broadly similar way, which complicates the interpretation of the results, as it becomes harder to know what information is necessary to predict each brain region. Structured variance partitioning provides a way to identify the information that is important in a voxel while taking into account the relationship between the layers. While we presented examples in this paper of a chain dependence between feature spaces (each layer is a function of the layer before it), this approach can be modified in future work to account for more complex relationships between feature spaces.

Traditional variance partitioning can be understood as an approach that approximates a conditional independence test. It is akin to asking if a brain area *Y* is related to a feature space *X* given another feature space *Z* (i.e., is *Y* ⊥ *X*|*Z*). Testing for conditional independence with a continuous *Z* is a particularly hard problem [70]. Variance partitioning approximates a solution for this problem, in a setting in which both *X* and *Z* are multivariate, by estimating the unique variance predicted by *X*. In that frame of mind, structured variance partitioning is akin to running a series of conditional independence tests while imposing constraints to reduce the complexity of the hypothesis space and improve interpretability. Variance partitioning is also similar to variable importance measures such as semi-partial correlation, which estimate the unique variance explained by one variable and not the others. Variance partitioning extends this type of variable importance measures where the variables are univariate to the setting where the variables are multivariate (i.e. the variables are feature spaces).

We have stated that because stacking *k* feature spaces results in our setting in models that are at least as good as models that use subsets of the *k* feature spaces, stacking is a good tool for structured variance partitioning (and variance partitioning in general). This is because it allows for more intuitive results (adding feature spaces will not result in a lower performance, and thus the unique variance explained will not be negative), and allows us to compute the set operations that are described in the methods. However, structured variance partitioning (and variance partitioning in general) can also be used with other ensemble learning methods for fMRI, as long as they combine feature space effectively and result in equal or greater preformance. Banded ridge [21, 22] for instance can be used as an estimation method in conjuction with structured variance partitioning.

We release with this paper a Python package to allow other researchers to use our method, which can be found at https://github.com/brainML/stacking. We hope that as data from naturalistic brain experiments become more accessible, there will be more progress in building encoding models with multiple feature spaces. Future work can provide better encoding performance and interpretability by considering nonlinear models, including comprehensive priors that include relationships between voxels, and further investigating improved interpretation methods.

## Acknowledgments

This research was supported in part by start-up funds in the Machine Learning Department at Carnegie Mellon University, and by the National Institute On Deafness And Other Communication Disorders of the National Institutes of Health under Award Number R01DC020088. The content is solely the responsibility of the authors and does not necessarily represent the official views of the National Institutes of Health.

## Supplementary Materials

**Supplementary Figure S1:**
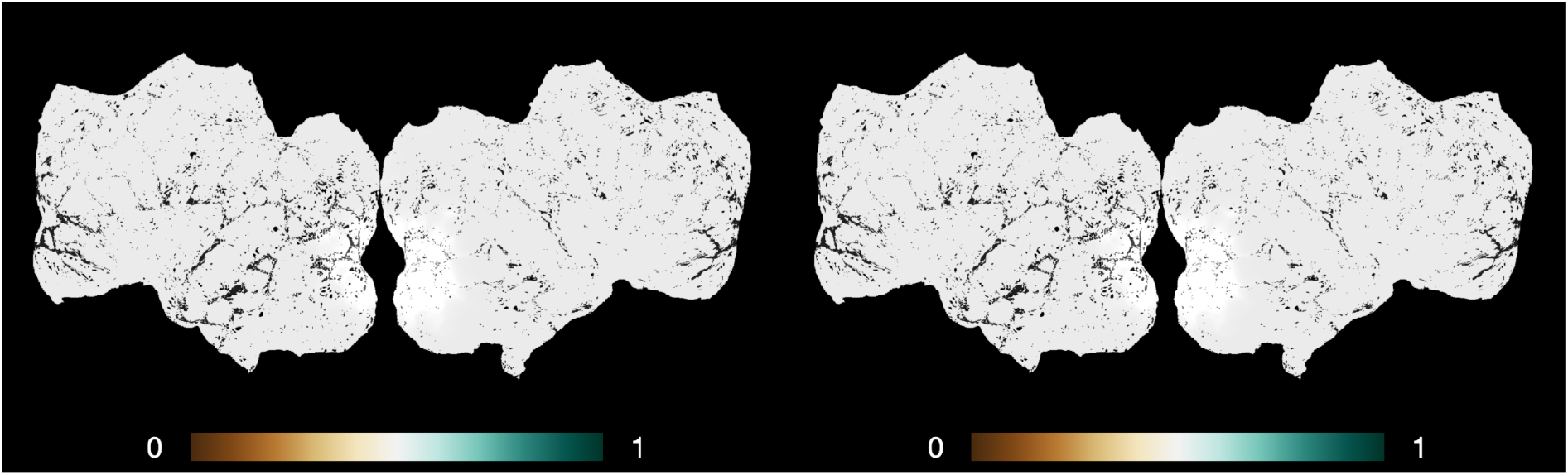
Stacking weights for two identical feature spaces shown on a participant’s flattened surface. The left map shows the weights for the first feature space and the right map shows the weights for the second feature space. We observe that the two figures are exactly the same, and every weight is 0.5. This result shows that our stacking method gives identical feature spaces the same weight. (Black pixels in both figures are missing voxels in the brain map.)

**Supplementary Figure S2:**
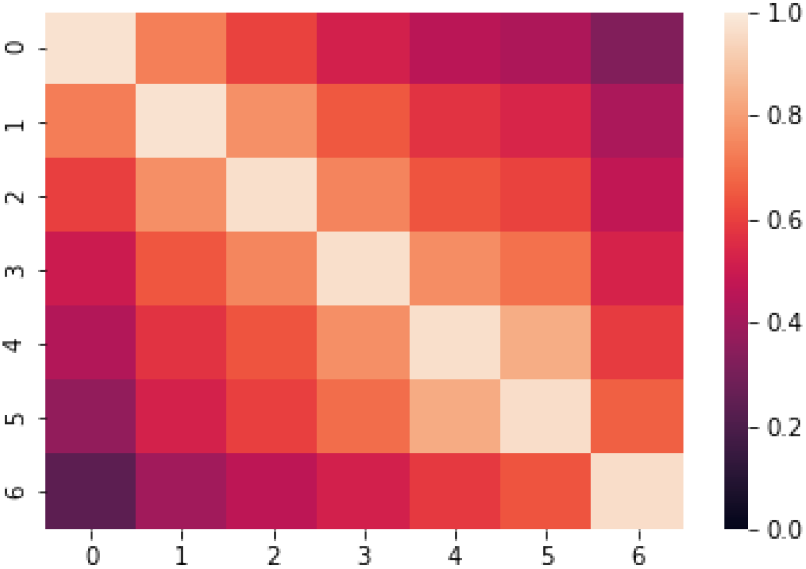
Similarity between the different layers of AlexNet, computed over the NSD images.

**Supplementary Figure S3:**
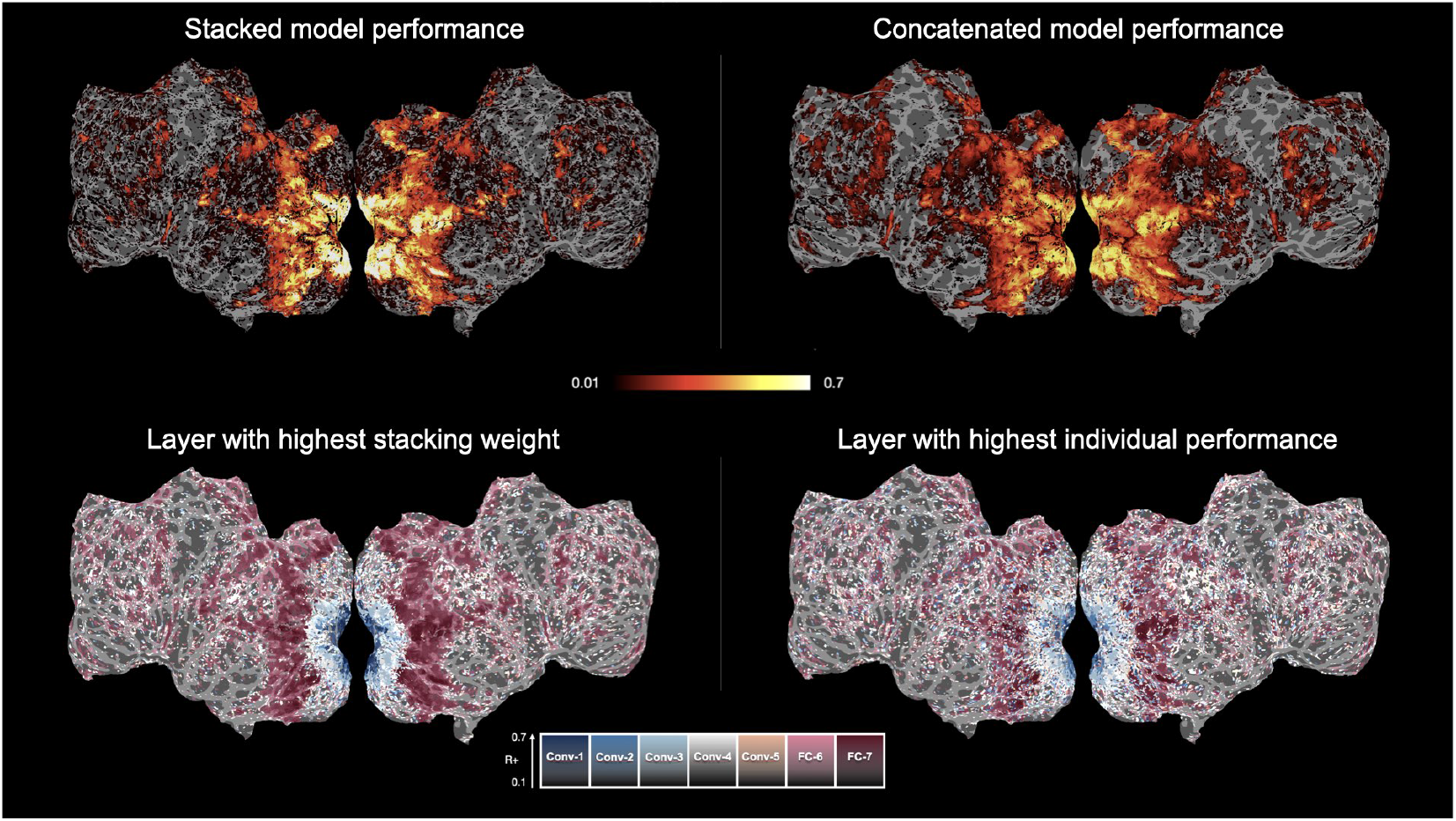
[Top] Stacking and concatenation prediction performance for subject S1. [Bottom] Feature attribution maps using the stacking weights (Left) and the layer with maximum performance (Right).

**Supplementary Figure S4:**
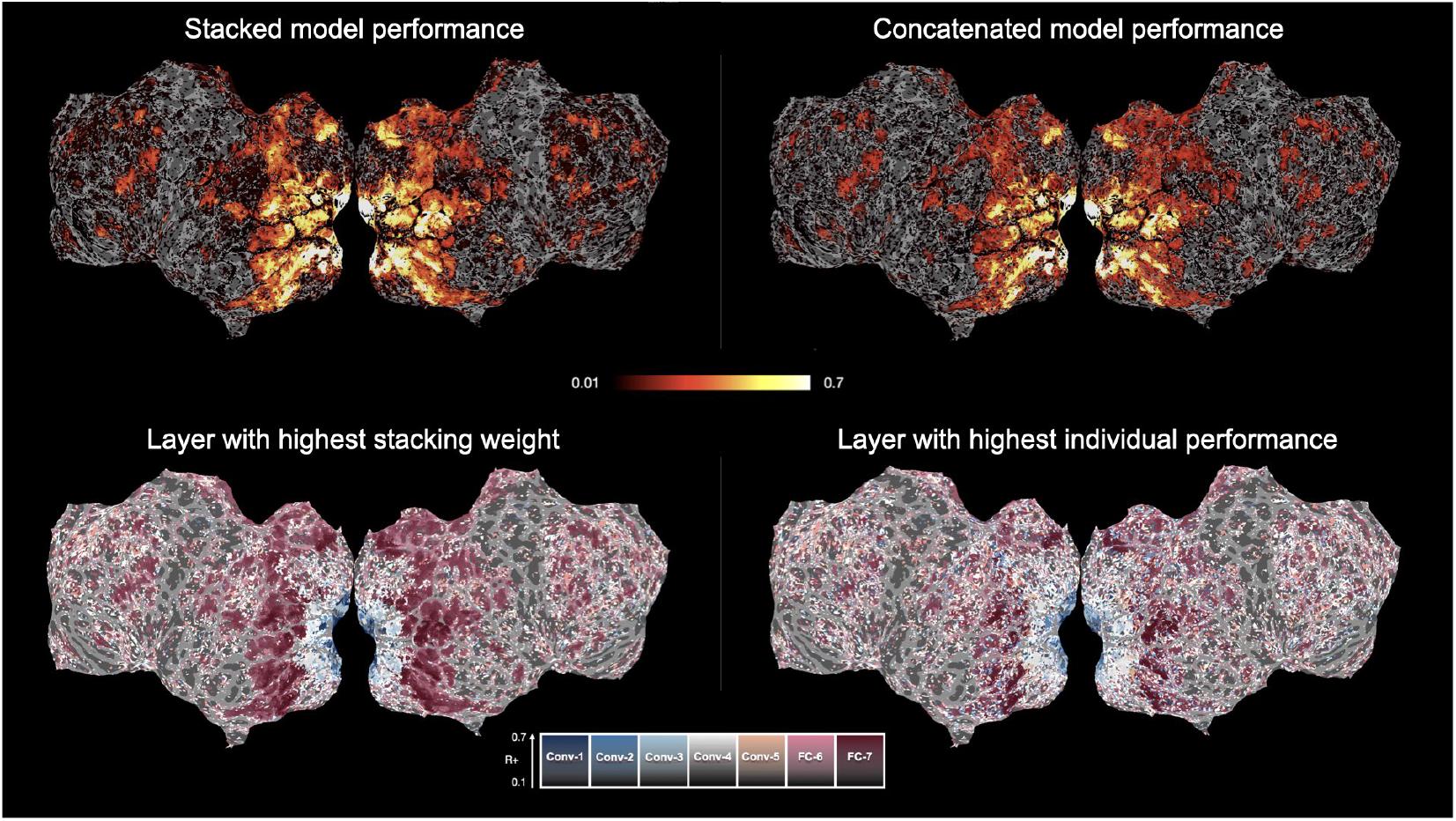
[Top] Stacking and concatenation prediction performance for subject S2. [Bottom] Feature attribution maps using the stacking weights (Left) and the layer with maximum performance (Right).

**Supplementary Figure S5:**
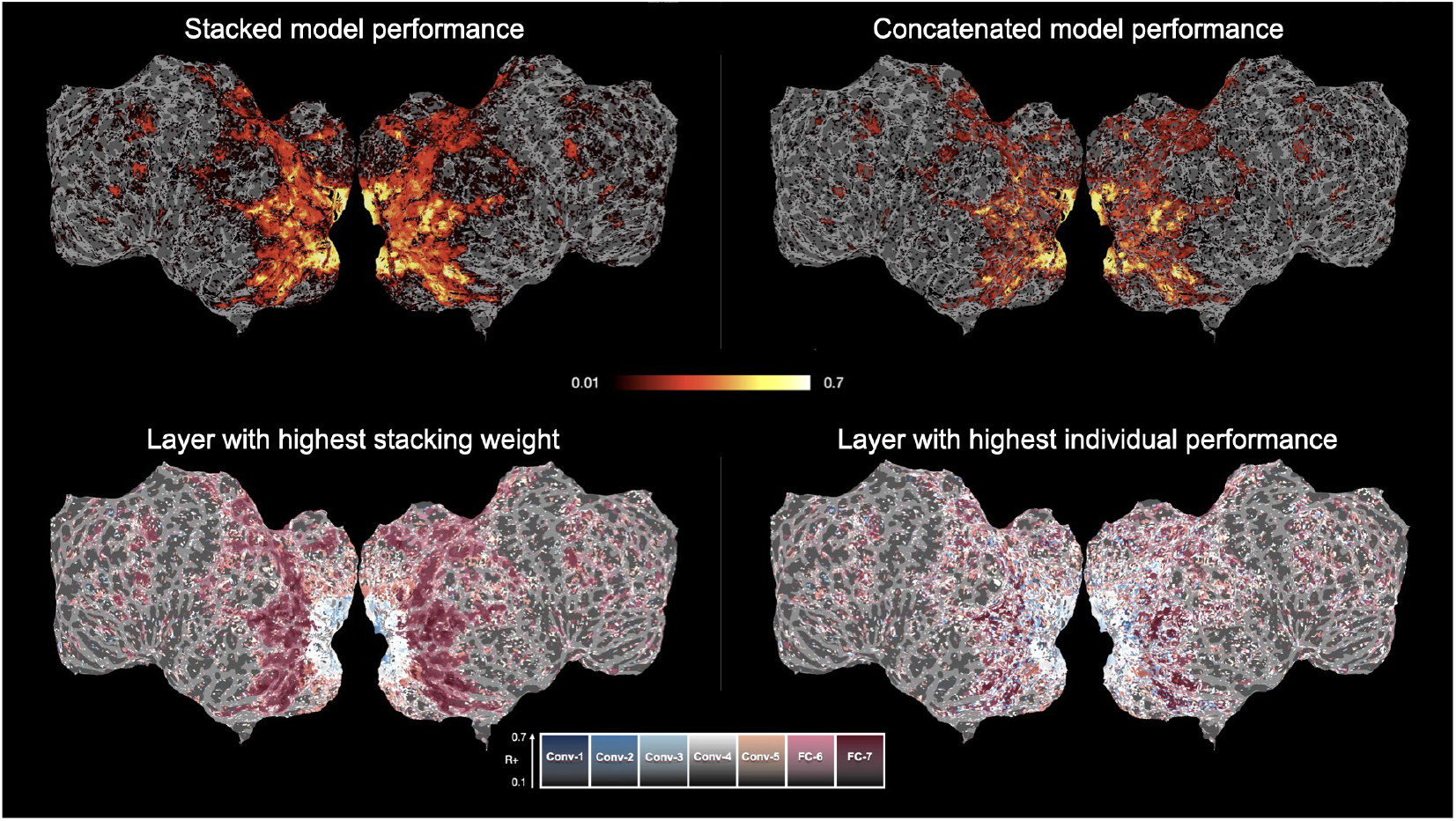
[Top] Stacking and concatenation prediction performance for subject S3. [Bottom] Feature attribution maps using the stacking weights (Left) and the layer with maximum performance (Right).

**Supplementary Figure S6:**
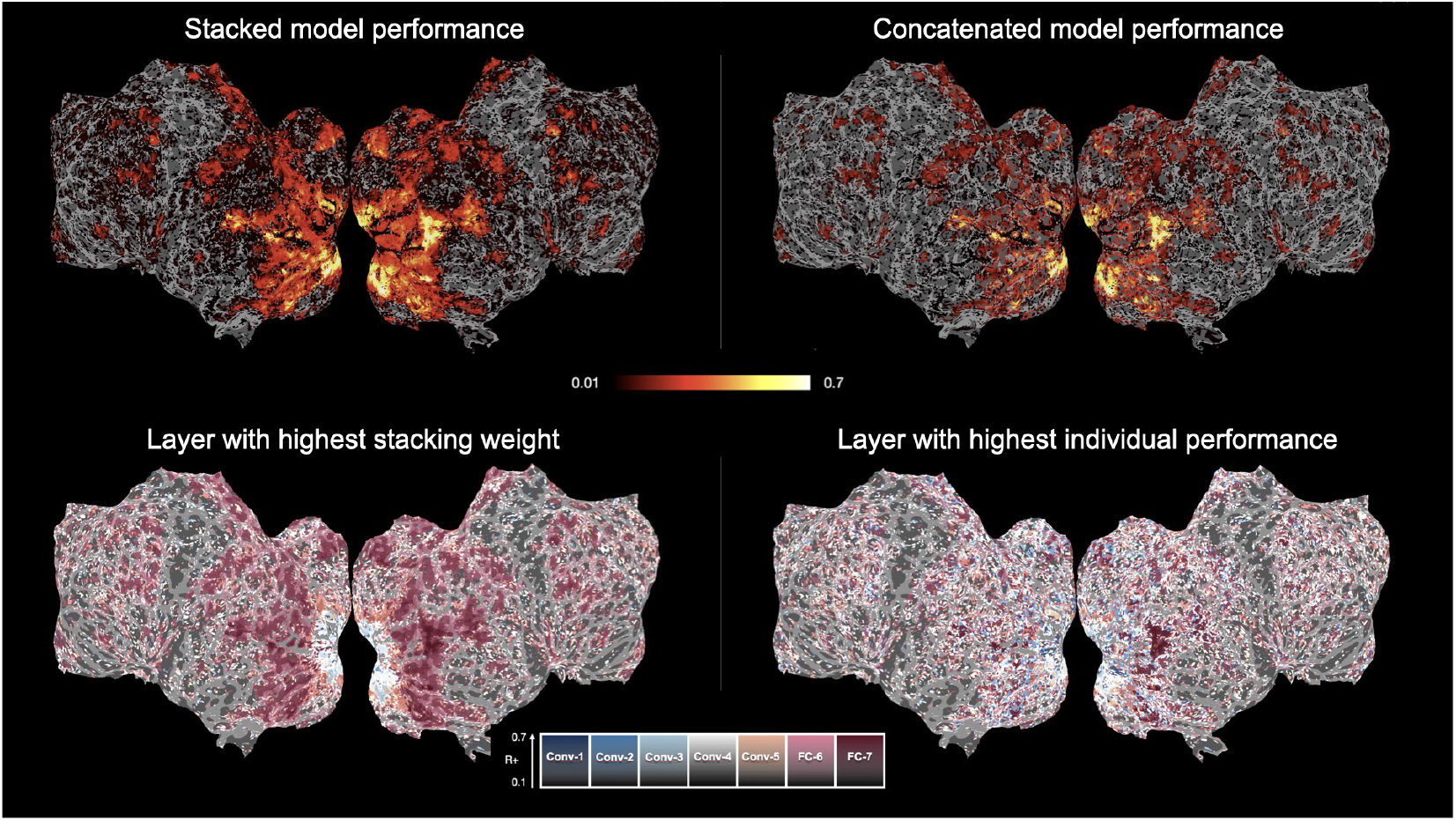
[Top] Stacking and concatenation prediction performance for subject S4. [Bottom] Feature attribution maps using the stacking weights (Left) and the layer with maximum performance (Right).

**Supplementary Figure S7:**
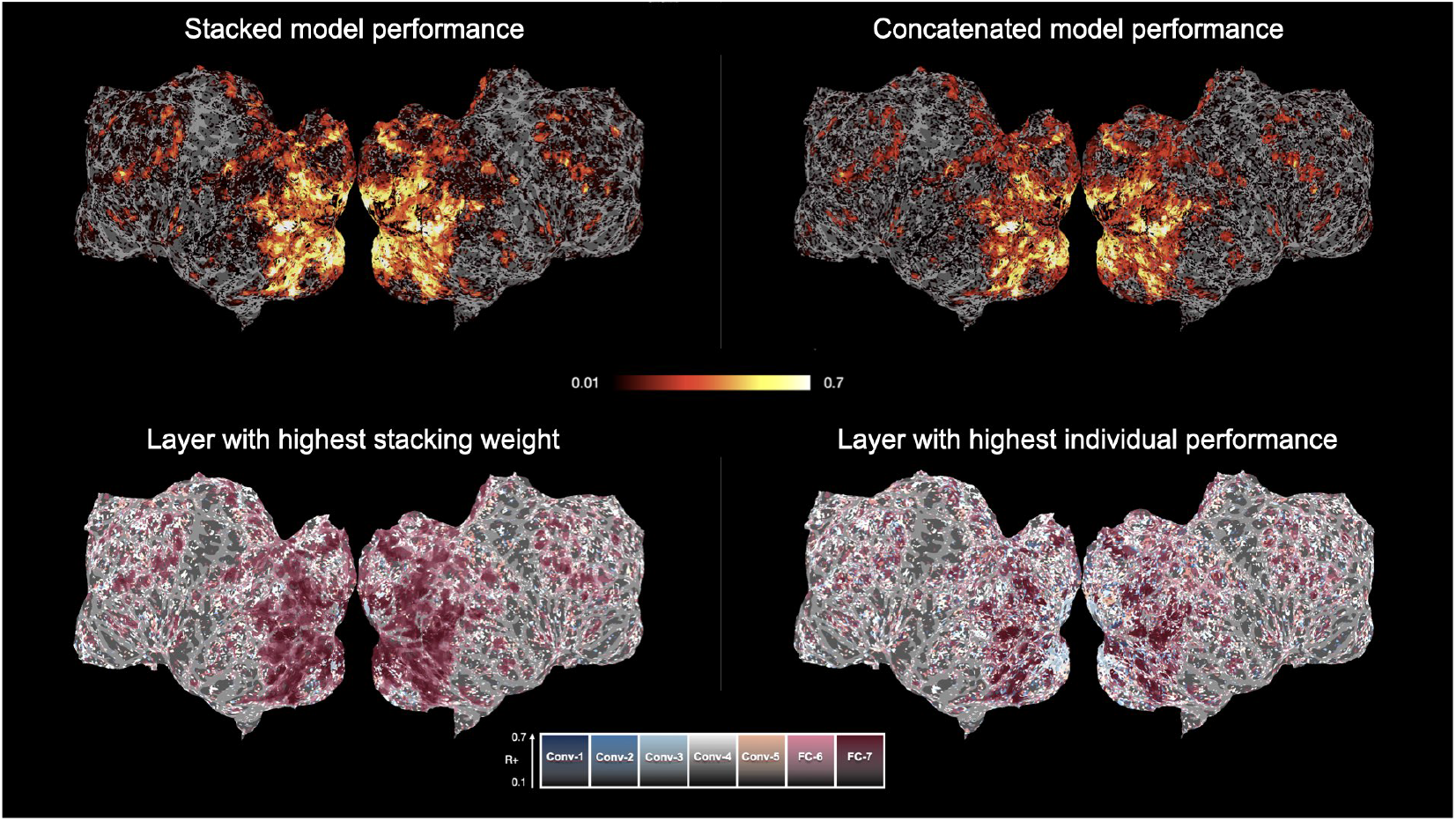
[Top] Stacking and concatenation prediction performance for subject S5. [Bottom] Feature attribution maps using the stacking weights (Left) and the layer with maximum performance (Right).

**Supplementary Figure S8:**
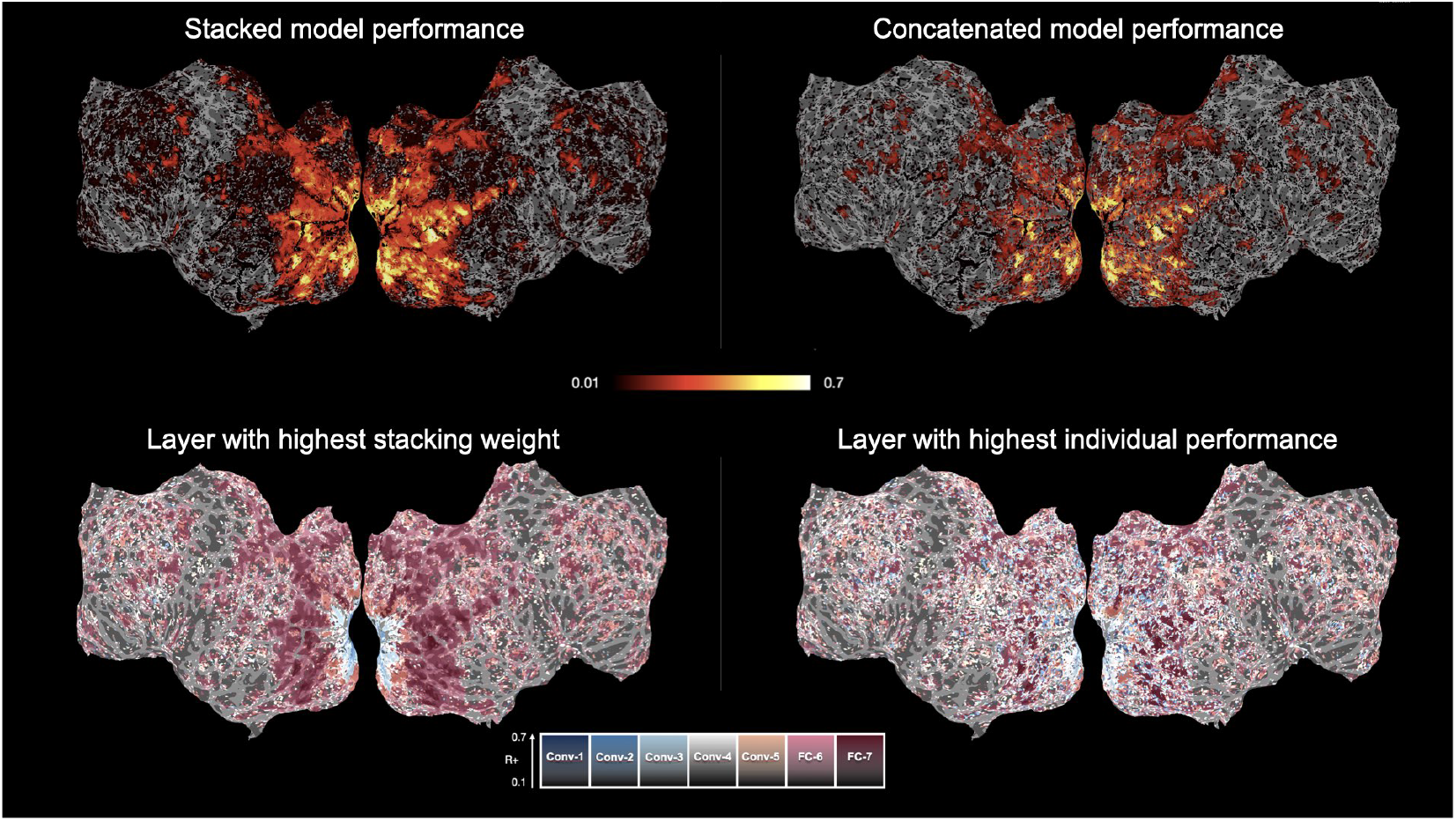
[Top] Stacking and concatenation prediction performance for subject S6. [Bottom] Feature attribution maps using the stacking weights (Left) and the layer with maximum performance (Right).

**Supplementary Figure S9:**
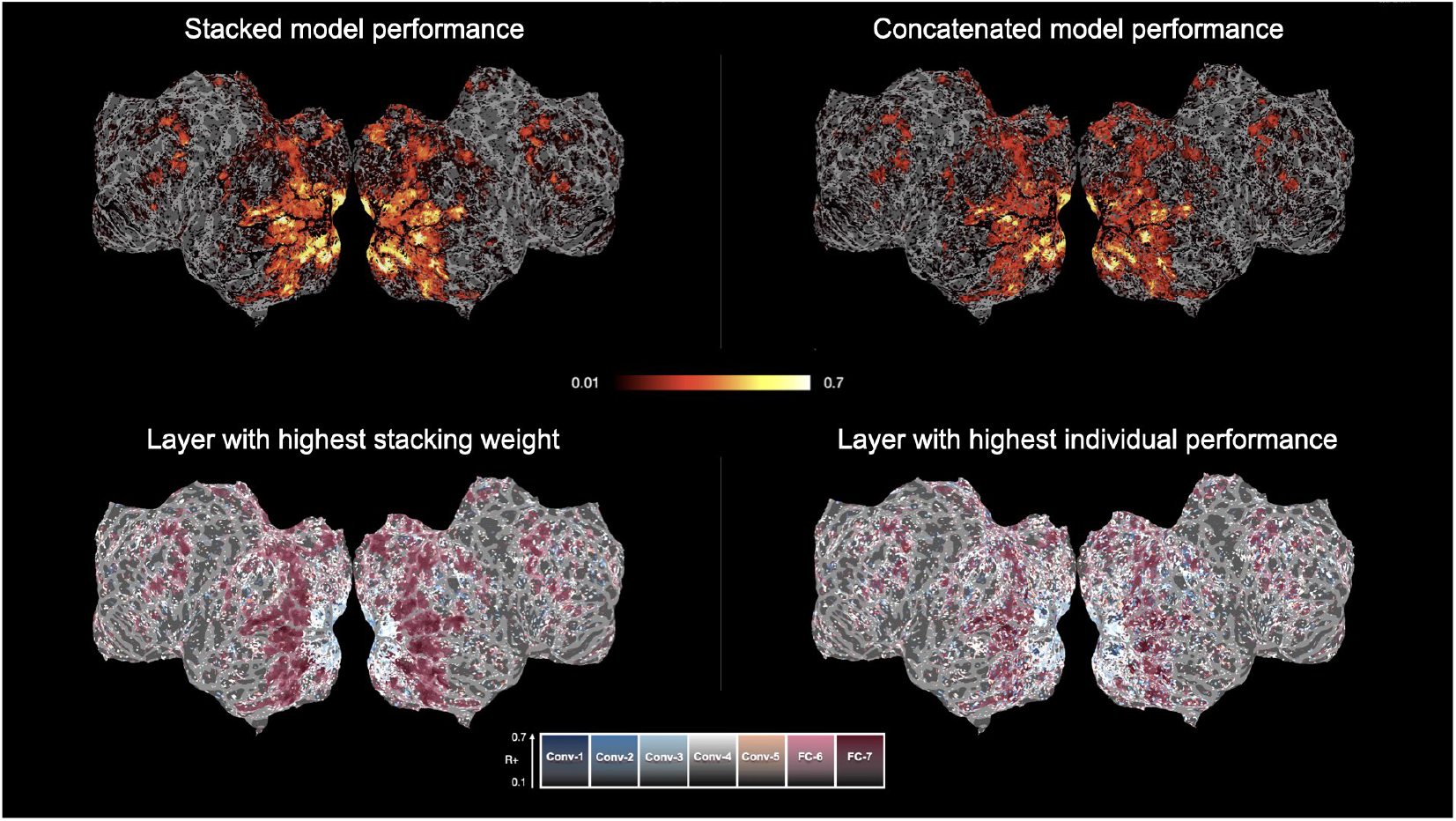
[Top] Stacking and concatenation prediction performance for subject S7. [Bottom] Feature attribution maps using the stacking weights (Left) and the layer with maximum performance (Right).

**Supplementary Figure S10:**
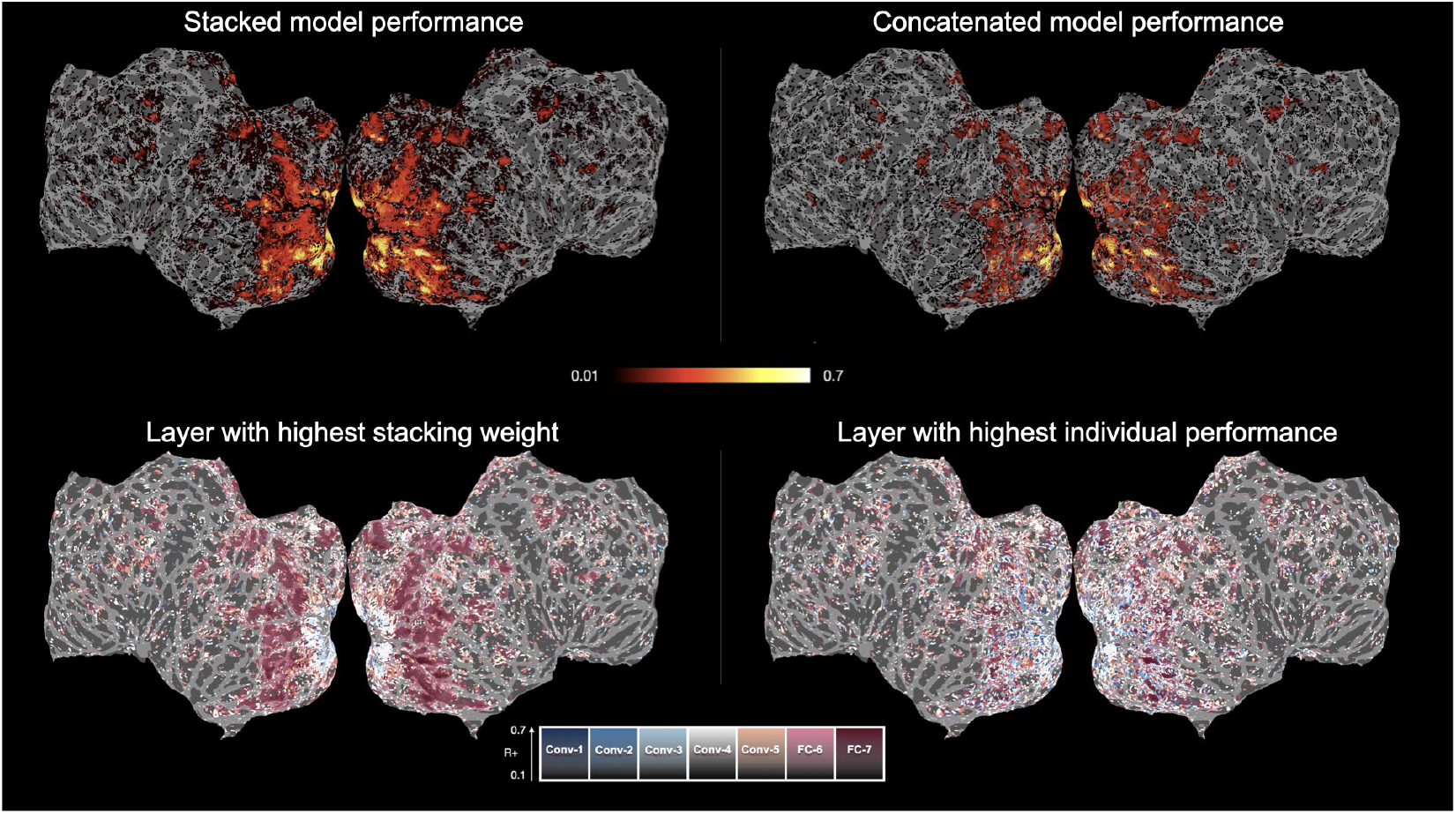
[Top] Stacking and concatenation prediction performance for subject S8. [Bottom] Feature attribution maps using the stacking weights (Left) and the layer with maximum performance (Right).

**Supplementary Figure S11:**
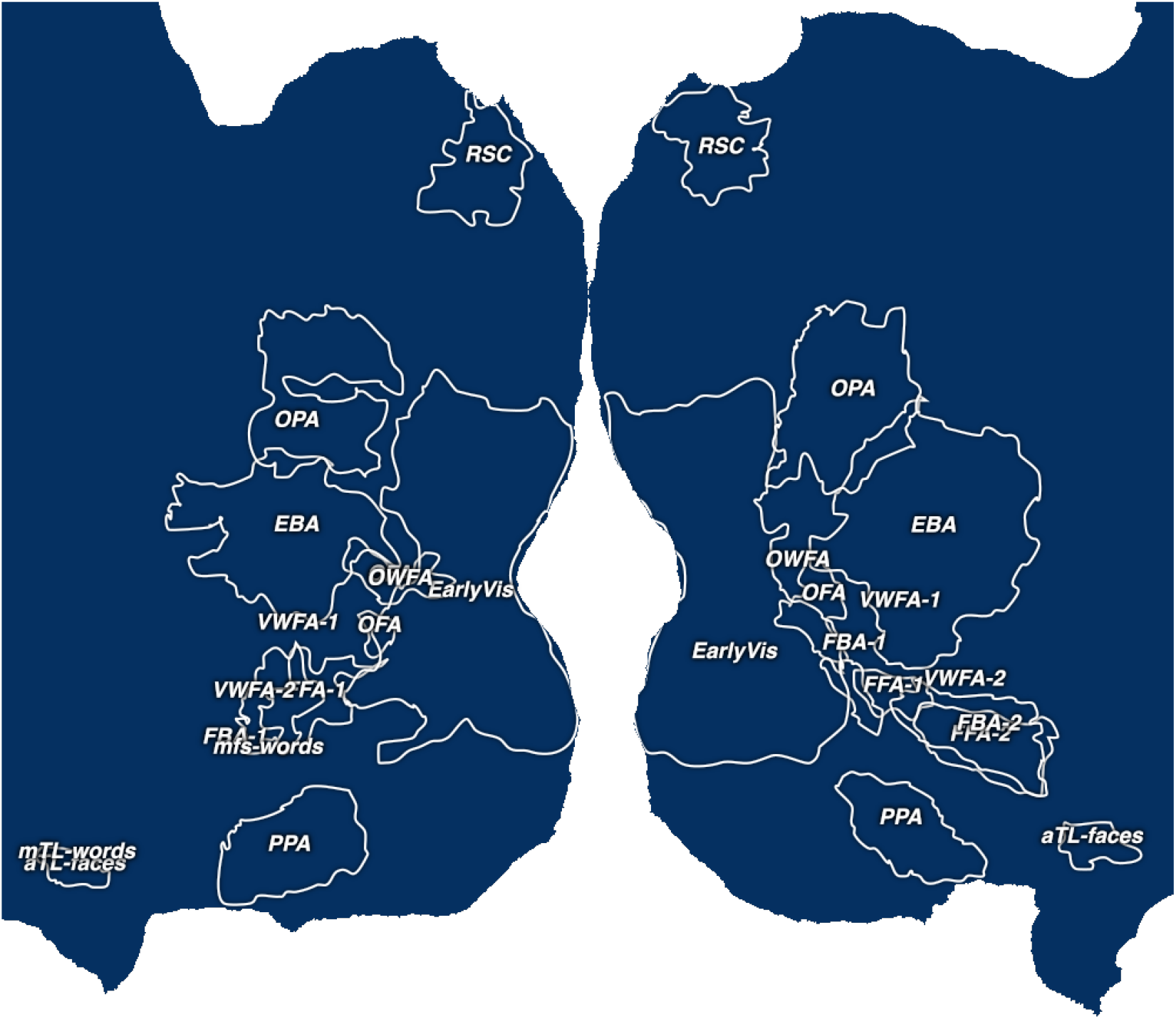
Flatmap representation of functional ROIs available in the NSD dataset for subject S1, obtained using the fLoc localizer [71].

**Supplementary Figure S12:**
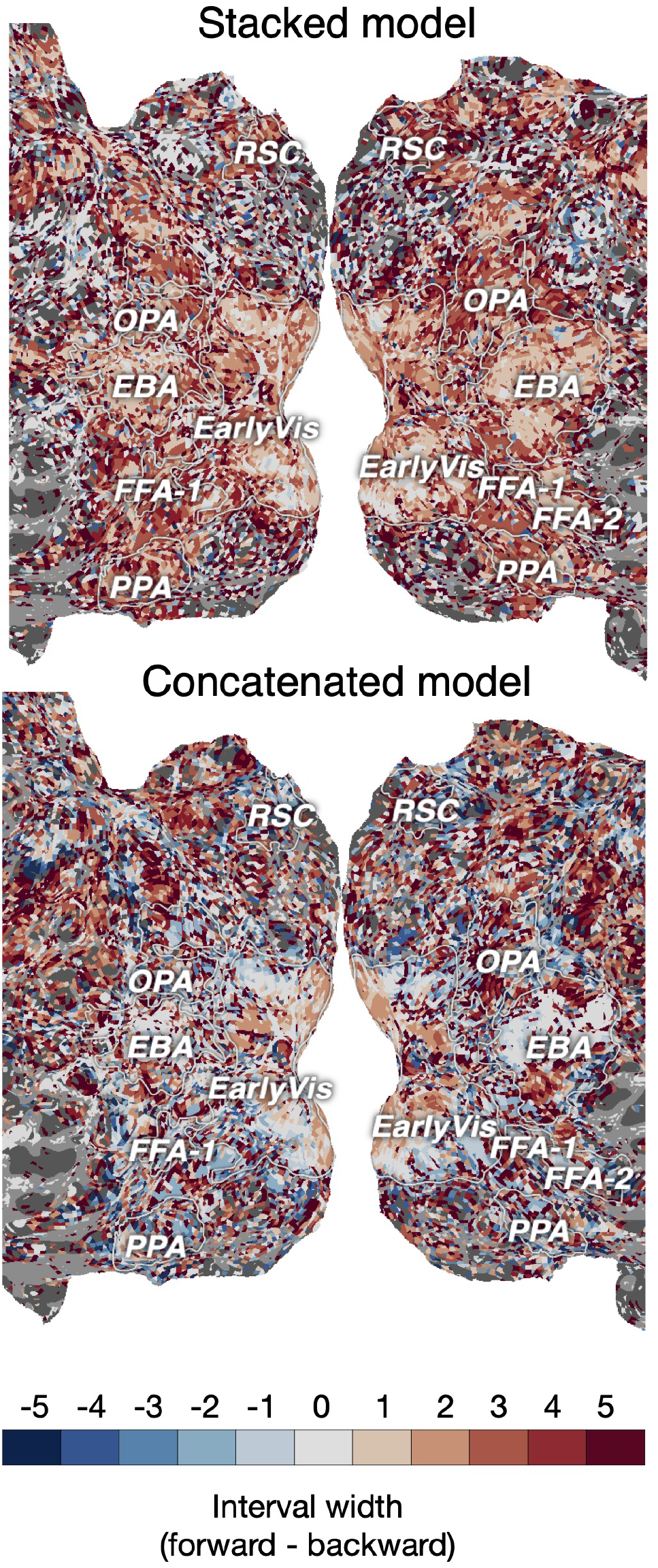
Cropped flatmap representation of the width of the interval between the layer obtained using the forward direction and the backward direction of structured variance partitioning, for subject S1.

## Notes

https://github.com/brainML/Stacking_Basics

### Competing Interest Statement

The authors have declared no competing interest.

